# Segmentally duplicated regulatory elements undergo human-specific rewiring

**DOI:** 10.64898/2025.12.03.692127

**Authors:** Seth Weaver, Craig B. Lowe

## Abstract

Gene regulatory innovation underlies many phenotypic transitions. Transposable elements are an established mechanism for creating families of cis-acting elements with shared sequence features and the potential to establish co-regulatory networks. To understand additional mechanisms by which co-regulatory networks form, we define families of noncoding elements based on sequence similarity and cell type-specific activity. We apply this analysis framework to the human telomere-to-telomere genome assembly and embryonic stem cell chromatin accessibility data. We identify segmental duplications as the major mechanism establishing these families, creating over one thousand networks of elements with open chromatin in embryonic stem cells. We functionally validate a subset of these networks as families of regulatory elements with STARR-seq and identify their target genes with CRISPRi in embryonic stem cells. Following segmental duplication, we find that regulatory elements at times maintain their relationship to target genes, and at times rewire to form novel connections. During this rewiring, we observe proximal-acting elements gaining the ability to regulate distally-located genes and observe transcriptional enhancers rewiring to regulate genes present at the locus outside the segmental duplication. Many of these rewiring events are human specific. Finally, we find that segmental duplications have made outsized contributions to expanding regulatory element families functioning in immune cell types and specific brain regions, including the posterior cingulate gyrus. We speculate that placing regulatory elements in new genomic contexts primes regulatory elements for neofunctionalization, and that regulatory rewiring after segmental duplication was a common mechanism underlying gene regulatory change during human evolution.

## Introduction

Coordinated gene expression is vital to organisms across all clades of life. In the case of multicellular organisms, sets of co-expressed genes are often distributed across the genome (Michalak, 2008; Razin et al., 2021). Coordinating the co-regulation of genes across the genome therefore requires dispersed *cis*-regulatory elements to act in concert, turning on or off their target genes at the same time (Britten and Kohne, 1968; Britten and Davidson, 1971). In order to accomplish this, these *cis*-acting regulatory elements likely share transcription factor binding sites, enabling them to act consistently in response to a change in the *trans*-regulatory environment (Yu et al., 2003; Allocco et al., 2004; Feschotte, 2008; Marco et al., 2009). These shared binding sites likely lead to DNA sequence similarities between regulatory elements that coordinate expression.

A well understood example of gene co-regulation in development is the coordinated expression of the *α*-like and *β*-like globin gene clusters, which reside on different chromosomes in tetrapods (Philipsen and Hardison, 2018). The coordination of expression across the two gene clusters is important for both maintaining the proper stoichiometric ratios of hemoglobin subunits and the transition between fetal and adult hemoglobin (Hardison, 2012; Sankaran and Orkin, 2013). This is accomplished by shared transcription factor binding sites, including GATA1/2 and TAL1, within the cis-regulatory elements at each cluster (Soler et al., 2010). While gene co-expression networks are known to be important, the mutational mechanisms by which they form are not fully understood.

One established mechanism for creating gene co-regulation is transposon insertions. Dating back to 1950, Barbra McClintock identified transposable elements (TEs) as “controlling elements,” due to their ability to control the expression of nearby genes (McClintock, 1950). Decades later, in the genomics era, interest was renewed in studying TEs in the context of forming gene regulatory networks (Bejerano et al., 2006; Thornburg et al., 2006; Polak and Domany, 2006; Lowe et al., 2007; Feschotte, 2008; Sundaram et al., 2014; Chuong et al., 2017; Ali et al., 2021; Oomen and Torres-Padilla, 2024). This body of work has elucidated that TE-derived regulatory elements may be widespread in their contributions to gene co-expression networks.

More recently, TEs have been studied and functionally validated for their contributions to immune responsive gene regulatory networks in mammalian systems (Grandi and Tramontano, 2018; Srinivasachar Badarinarayan and Sauter, 2021). In mice, B2 SINE elements contain STAT1 binding sites and act as type II interferon-inducible enhancers (Horton et al., 2023). In humans, a primate-specific endogenous retro viral family, MER41, was also found to shape interferon-response gene regulatory networks (Chuong et al., 2016). This recent work highlights both the contribution of TEs in establishing gene co-expression networks and the importance of studying lineage-specific duplication events for understanding the evolution of gene co-regulatory networks (Teichmann and Babu, 2002).

Along with detailed case studies of TEs contributing to the co-regulation of genes, there was a more general genome-wide screen to first infer the location of regulatory elements using cross-species conservation and then use DNA sequence similarity to group them into families (Bejerano et al., 2004). A strength of this approach is that it can identify putative regulatory elements that share significant sequence identity and are therefore likely to share binding sites and functions, regardless of the cell type in which they are active, or the mutational mechanisms that created them. Several of these families were the result of ancient transposons spreading regulatory elements (Bejerano et al., 2006; Xie et al., 2006), but how the majority of these families originated is not known. We hypothesized that segmental duplications (SDs) may also contribute to building families of regulatory elements. SDs are already well-known for duplicating and dispersing protein-coding genes (Ohno, 1970; Sharp et al., 2005; Dennis and Eichler, 2016; Dougherty et al., 2018; Fiddes et al., 2018b) and recent discoveries have emphasized the large extent to which SDs have shaped, and continue to shape, our genomes. A key example with protein-coding genes is the expansion of the *SRGAP2* gene family, which contributed to human-specific phenotypes, such as our larger neocortex and greater synaptic density (Dennis et al., 2012; Schmidt et al., 2019). The SDs affecting the *SRGAP2* gene family are not isolated incidents with ∼7% of the human genome recently originating from SD events (Vollger et al., 2022). Additionally, SDs account for roughly 1100 gene copy number variants per diploid genome in human populations (Jeong et al., 2025), indicating that SDs have made widespread contributions to both what make the human genome unique compared to other species (Blekhman et al., 2009; Ju et al., 2016; Nuttle et al., 2016), as well as what makes individual humans unique when compare to each other (Cooper et al., 2011).

While the historical focus has been on how SDs affect protein-coding regions, the ability of SDs to influence gene regulation is beginning to be understood. Originally, this was studied in the context of gene duplication as a mechanism for upregulating protein expression, such as amylase copy number in human populations (Perry et al., 2007; Yilmaz et al., 2024). It is now being increasingly appreciated that large SDs carry diverse regulatory elements that affect gene regulation over long distances (Wu et al., 2017; Torres et al., 2024). As an example, the tandem duplication of a distal enhancer for SOX9, but not SOX9 itself, leads to an upregulation of SOX9, causing sex reversal in humans (Kim et al., 2015). When the duplication event is larger, and contains an insulator along with the enhancer, the duplicated enhancer does not contact SOX9, resulting in no phenotypic change (Franke et al., 2016). Even longer tandem duplications have the duplicated SOX9 enhancer forming a novel contact with KCNJ2, resulting in Cook’s Syndrome. These examples illustrate that gene regulatory elements remain active post-duplication and may continue to regulate genes they were duplicated with, “rewire” to contact a new gene, or become “orphaned” and not regulate any genes despite having enhancer activity in isolation.

To address this question of how families of regulatory elements originate, we revisited the analysis of noncoding element families with modern data sets and experimental techniques. We perform sequence-similarity clustering within the telomere-to-telomere human reference genome (Nurk et al., 2022) to identify families of sequences where their paralogy is not solely due to transposon sequences. We combine this paralogy map with epigenomic signals to define cell type-specific noncoding element families that are strong candidates for regulating genes. We identify SDs as the primary genomic mechanism for creating and expanding families of putative regulatory elements. We use both self-transcribing active regulatory region sequencing (STARR-seq) (Arnold et al., 2013) and CRISPR-interference (CRISPRi) screening (Thakore et al., 2015) to functionally validate members of these noncoding element families for enhancer activity *in vitro* and identify their target genes. We uncover that enhancers may be “masked” or “orphaned” by their genomic context, but can regain the ability to influence the transcription of a target gene upon duplication into a new locus. We also observe distal enhancers rewiring to new target genes upon duplication, as well as a fluidity in which promoter elements appear to easily gain distal gene regulatory function following their duplication.

## Results

### Sequence clustering of noncoding elements with cell type specificity

We first propose an analysis framework to define clusters of noncoding elements based on DNA sequence similarity and epigenetic signals (Fig. 1A). Briefly, the first step is to generate a genome-wide self-alignment to identify regions of paralogy that are not solely based on transposon sequences (see Methods). Historically, it was difficult to properly assess paralogy in genome assemblies because highly divergent haplotypes would at times be assembled as paralogs, and highly similar paralogs would often be collapsed as if they were divergent haplotypes. To address these issues, we used the telomere-to-telomere human genome assembly (Nurk et al., 2022). The second step is to define a set of elements that potentially have a noncoding function. While this was previously done by focusing on cross-species DNA sequence conservation outside of protein-coding exons, there are now a large number of biochemical data sets that can identify elements with noncoding functions. This paradigm shift has been important because these datasets include recently evolved regulatory elements and allow us to know the tissue and cell type of activity. While many biochemical marks can help identify putative regulatory elements, we utilize chromatin accessibility in this study due to its ability to capture a broad range of functional elements. For simplicity, we refer to the putatively functional elements identified by our epigenetic data as “open,” but emphasize that this approach could utilize assays beyond chromatin accessibility. Along with using the self-alignment to identify homology between open elements, we also identify regions of high sequence similarity between open elements and regions that do not show signs of open chromatin.

**Fig. 1.**
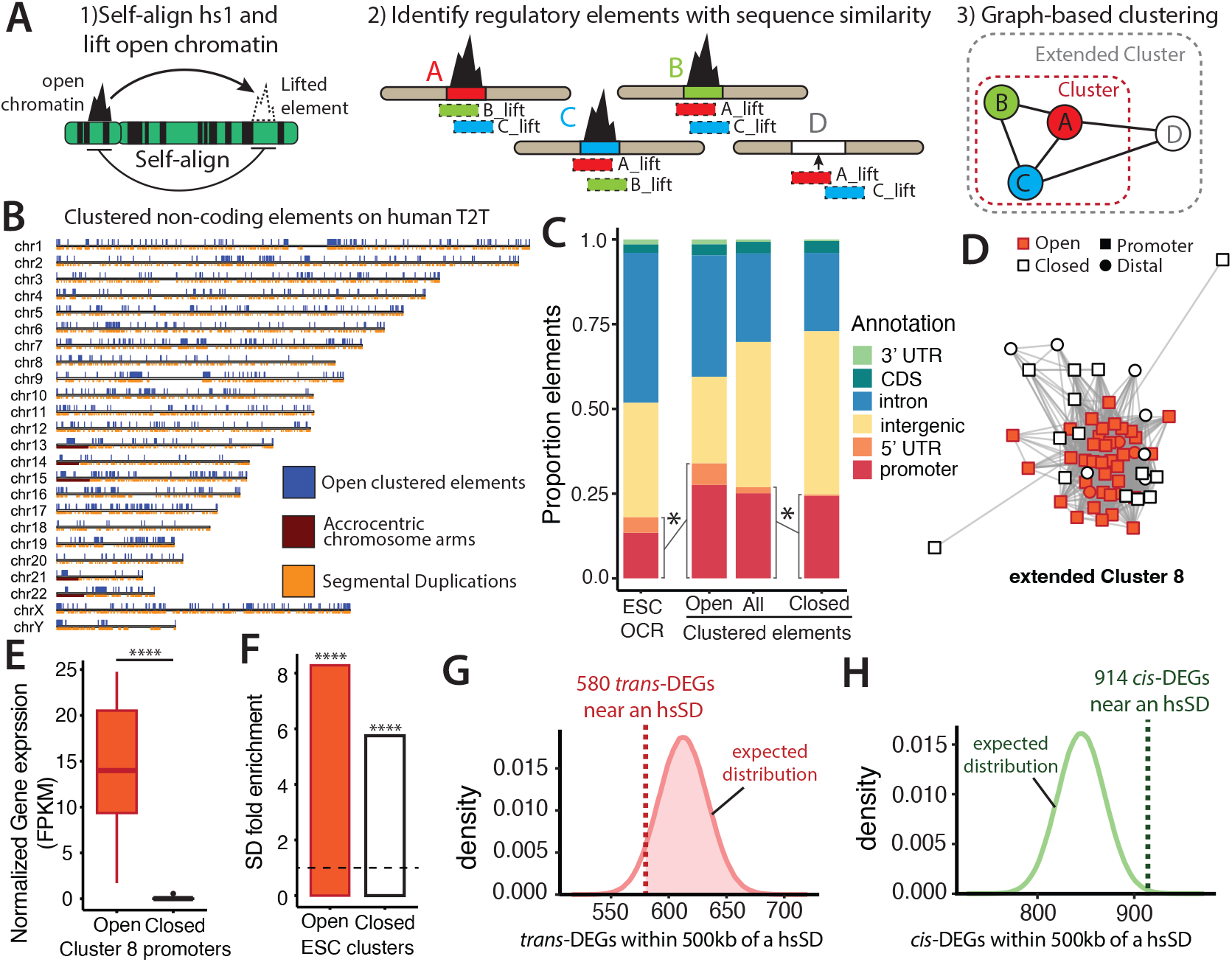
Clustered noncoding elements in the human genome. **(A)** Overview of method for cell type-specific clustering of noncoding elements. **(B)** Location of embryonic stem cell (ESC) clustered elements, segmental duplications and acrocentric chromosome arms in the human T2T reference genome. **(C)** Comparison of genomic annotations of all ESC open chromatin regions (OCRs) to the subset that appear in clusters, and the comparison of all clustered elements to closed clustered elements (binomial, *: p *<* 0.0001). **(D)** Extended Cluster 8 visualized with edge lengths corresponding to percent divergence between noncoding element nodes. **(E)** Normalized expression of genes with open and closed Cluster 8 promoters (t-test, ****: p *<* 0.0001). **(F)** Fold enrichment of segmental duplications (SDs) for overlapping open and closed ESC clustered elements (binomial, ****: p *<* 0.0001). **(G**,**H)** The number of human-chimpanzee differentially expressed genes due to *trans-* and *cis-*regulatory changes within 500kb of a human-specific SD (hsSD). These observed counts are compared to null distributions corresponding to randomly sampling from all genes in the genome one million times.

We define this putatively non-functional (in the given cell type) set of elements as “closed homologous elements;” for simplicity, we often refer to these as “closed elements.” In total, we use the sets of open and closed elements to create graph-based clusters of noncoding elements with shared sequence features. We define “clustered noncoding elements” (“clusters”) as connected components within a graph where nodes are open elements (based on epigenetic signal) and edges represent significant DNA sequence similarity. “Extended clusters” are connected components that contain open elements as well as nodes representing closed elements. This computational framework is able to identify cell type-specific clusters of noncoding elements with DNA sequence and epigenetic features that potentially underlie gene co-regulatory networks.

We applied this analysis framework to H9 embryonic stem cells (ESCs), a biologically relevant and experimentally tractable cell type. As a proxy for gene regulatory activity, we re-aligned ATAC-seq reads from H9 ESCs (Liu et al., 2017) to the T2T human genome (see Methods). This resulted in 126,217 open chromatin peaks, which are each represented by a node in a graph. We used the self alignment to add edges to the graph (see Methods), which resulted in 1336 clusters containing at least 2 elements open in ESCs (Table S1). These clusters contain a total of 4298 open chromatin elements, which are located across all chromosomes (Fig. 1B) and have a mean length of 325 bp (median = 250 bp) (Figure S1A). The ESC clusters range in size from 2 to 92 noncoding element nodes with an average size of 3.29 elements (median = 2).

When extended ESC clusters are considered, an additional 13,944 closed homologous elements are added to the existing clusters, with a mean extended cluster size of 13.73 (median = 3) (Figure S1B). There exists considerable variability in the open/closed element composition of extended clusters, evidenced by the lack of correlation between the number of open and closed elements per cluster (*R*^2^ = 0.017) (Figure S1C). We note that although here we define function on the basis of ESC epigenetic signal, ESC closed elements may have chromatin accessibility in other cell types. We overlapped ESC closed elements with epigenetic data from 154 different ENCODE cell types, and observed 68% of ESC closed elements are open in at least one other cell type (Figure S1D). In summary, we identified over 1000 clusters of noncoding elements in ESCs that have the shared sequence components and epigenetic signals to potentially establish gene co-regulatory networks.

### Clusters are biased towards proximal elements

To understand how clusters of noncoding elements could be influencing gene co-regulation, we more finely categorized their potential function. One-third of the elements in ESC clusters are likely to be proximally acting, based on being located in promoters (within 2 kb upstream of a transcription start site) or 5’ UTRs, which is a 1.8-fold enrichment compared to all ESC open chromatin peaks (*p <* 10^−20^; Fig. 1C). The enrichment for proximal elements suggests that gene duplication is a major mechanism for the expansion of ESC clusters and that clustered promoter elements may continue to be open in the same cell types, co-regulating duplicated genes to increase expression, such as with amylase genes (Yilmaz et al., 2024). In contrast, the subset of closed elements in extended ESC clusters are mildly depleted for proximal elements compared to the the combined open and closed element set (Fig. 1C) (fold-depletion = 0.92, *p <* 10^−20^). The depletion of proximal elements in the closed subset is consistent with our previous result, suggesting that proximal elements are more likely than distal elements to remain open after a recent duplication.

Though promoter elements are enriched to appear in clusters (compared to distally acting elements), two-thirds of ESC clustered elements are likely to be distally acting. Unlike proximally acting elements where the target gene is easier to infer, it is difficult to predict how duplicated distally acting noncoding elements will act in their new location. Even experimental methods, such as Hi-C, often have a limited capacity to assign potential target genes to clustered elements because multi-mapping sequence reads are typically removed from the analysis (Zheng et al., 2019). We hypothesize that duplicated distal elements will fall into one of three categories: maintaining regulatory relationships with genes they were duplicated alongside, rewiring into the existing regulatory architecture at the new locus, or becoming masked/orphaned where they do not regulate a gene despite having the ability to do so in isolation.

### Reliability of epigenetic data in paralogous regions

Many functional genomic assays rely on being able to accurately map short sequencing reads back to the genome, which can be difficult when multiple regions have a high amount of sequence similarity. However, this issue may have been ameliorated over time by short sequencing technologies progressively offering longer reads and a T2T reference genome addressing inaccuracies in the assembly itself. To assess if our open chromatin data was accurately aligning to truly open elements, and not aligning to the closed chromatin subset, we analyzed extended Cluster 8 (Fig. 1D), a cluster of 56 elements where 49 are promoters. We assessed the agreement between the inference of promoter activity based on chromatin status versus the inference of promoter activity based on the expression of the associated transcript in RNA-seq. In extended Cluster 8, genes with open chromatin promoters had higher average expression in ESCs compared to genes with closed element promoters (t-test; *p <* 10^−11^) (Fig. 1E). This is not an extended Cluster 8 specific phenomenon: genome wide, genes with open clustered noncoding element promoters have higher average expression in ESCs than genes with closed homologous element promoters (Wilcoxon; *p <* 10^−11^) (Figure S1E). The elevated gene expression of open promoter genes is evidence that epigenetic signals can be reliably mapped to sequence families and that our analysis framework is properly segregating open and closed subsets of clustered noncoding elements.

### Sequence similarity predicts functional similarity

We observed open Cluster 8 elements congregating densely in the center of the graph with high sequence similarity to each other, while the closed elements were dispersed along the periphery (Fig. 1D). Confirming this observation, Cluster 8 open chromatin elements have a lower percent divergence to other open elements than to closed elements, or than closed elements have to each other (Figure S1F). We suggest that conserved sequence features within the open Cluster 8 elements are important for their function and that drifting too far from these conserved sequence features leads to the loss of gene expression in ESCs.

### Segmental duplications create noncoding clusters

We set out to determine the mutational mechanisms responsible for the creation of the observed clusters of noncoding elements in ESCs. Since SDs are the largest source of novel euchromatic sequence in human evolution (Dennis and Eichler, 2016; Vollger et al., 2022) and are well-established as a mechanism for creating clusters of protein-coding genes (Dennis et al., 2012; Fiddes et al., 2018b), we hypothesized that SDs may be creating and expanding noncoding element clusters. To test this hypothesis, we overlapped SDs annotated in the T2T human assembly (Vollger et al., 2022) with all open and closed elements. We identified that annotated SDs are enriched 8.3-fold for open clustered elements (*p <* 10^−10^) and 6.8-fold for closed homologous elements (*p <* 10^−10^) (Figure 1F). Out of 4298 ESC open elements, 3387 (82%) are contained within SDs, and 878 SDs have at least one ESC open element. Out of 1336 clusters, 1057 (79%) have at least one member in an annotated SD and 974 clusters (73%) have at least 75% of their members within SDs. Taken together, this analysis indicates that SDs are a key mutational mechanism for duplicating and dispersing noncoding elements throughout the human genome.

### Segmental duplications change the expression of neighboring genes

While SDs are important for creating and expanding clusters of noncoding elements, it remains unknown how these elements function and if SD-duplicated elements maintain previous functions in their new location. In the cases where the functional noncoding elements are duplicated alongside genes, the local regulatory architecture may be preserved with the duplicated element maintaining existing regulatory relationships internal to the SD. A second option is that the functional noncoding element may become masked/orphaned, maintaining regulatory potential, but without regulating a gene due to the chromatin landscape or insulating features in the genome. A third possibility is that duplicated functional elements may integrate into the existing gene regulatory architecture at the new locus. While there are case studies supporting the possibility of these three outcomes at a locus (Franke et al., 2016), the relative proportions remain unknown. We hypothesized that if it is common for SDs to influence the existing regulatory architecture of neighboring genes at a new locus, as opposed to being self contained or masked/orphaned, we would be able to detect an association between the lineage-specific expansion of clustered noncoding elements and the differential expression of neighboring genes. To test this hypothesis, we investigated if human lineage-specific SDs tend to be located next to genes that are differentially expressed between humans and chimpanzees. We generated a subset of SDs that are human-specific by removing any SDs within a syntenic alignment to Oxford Nanopore (ONT) long-read sequencing data from a chimpanzee (Yoo et al., 2025) and were therefore already present in the human-chimpanzee ancestor (see Methods). We analyzed allele-specific expression data from human-chimpanzee allotetraploid iPSCs (Gokhman et al., 2021; Pavlovic et al., 2022) and subsetted the differentially expressed genes into those affected by *cis* or *trans* regulatory changes. We found that while human-chimpanzee differentially expressed genes caused by *trans*-regulatory changes are not enriched for being located near human-specific SDs (*p* = 0.94) (Figure 1G), differentially expressed genes caused by *cis* changes are enriched within 500kb of human-specific SDs (*p* = 0.003) (Fig. 1H). The enrichment of differentially expressed genes caused by changes in *cis* to be located near human-specific SDs is consistent with SDs often modifying the local gene regulatory architecture upon insertion.

### Functional interrogation of segmentally duplicated noncoding elements with STARR-seq and CRISPRi

While there could be multiple explanations for human-specific SDs affecting the expression of nearby genes, such as disrupting existing regulatory interactions, we hypothesize that the clustered noncoding elements in SDs retain their function upon duplication, at times acting within the SD, and at times acting on genes outside the SD. While SDs can duplicate diverse classes of noncoding elements, we chose to focus on clustered noncoding elements that have biochemical marks of promoters and transcriptional enhancers. Additionally, we prioritized clusters that were expanded by human-specific SDs, and could potentially be contributing to aspects of human-specific biology (Vaill et al., 2023). To assay if the clustered noncoding elements identified in our screen have regulatory element activity *in vitro*, and identify the gene(s) they target, we performed two orthogonal functional assays in H9 ESCs.

First, we performed an episomal enhancer assay, self-transcribing active regulatory region sequencing (STARR-seq) (Arnold et al., 2013) to measure enhancer activity in 112 open and closed elements across 11 different noncoding element clusters in ESCs (Figure 2A). For each tested element, we quantified enhancer activity by relating the abundance of each test sequence in the transcriptome to the abundance of the test sequence in the plasmid library. We further calculated relative enhancer strength compared to the activity of 18 negative controls to identify test constructs with significant enhancer activity (*p <* 0.01) (see Methods). We performed three replicates and the enhancer activity across the replicates were well correlated (avg. R^2^ = 0.96) (Figure S2A). The strength of this assay is that it is highly quantitative and tests for the enhancer activity of a DNA sequence when isolated from the endogenous context of chromatin architecture and target gene(s).

**Fig. 2.**
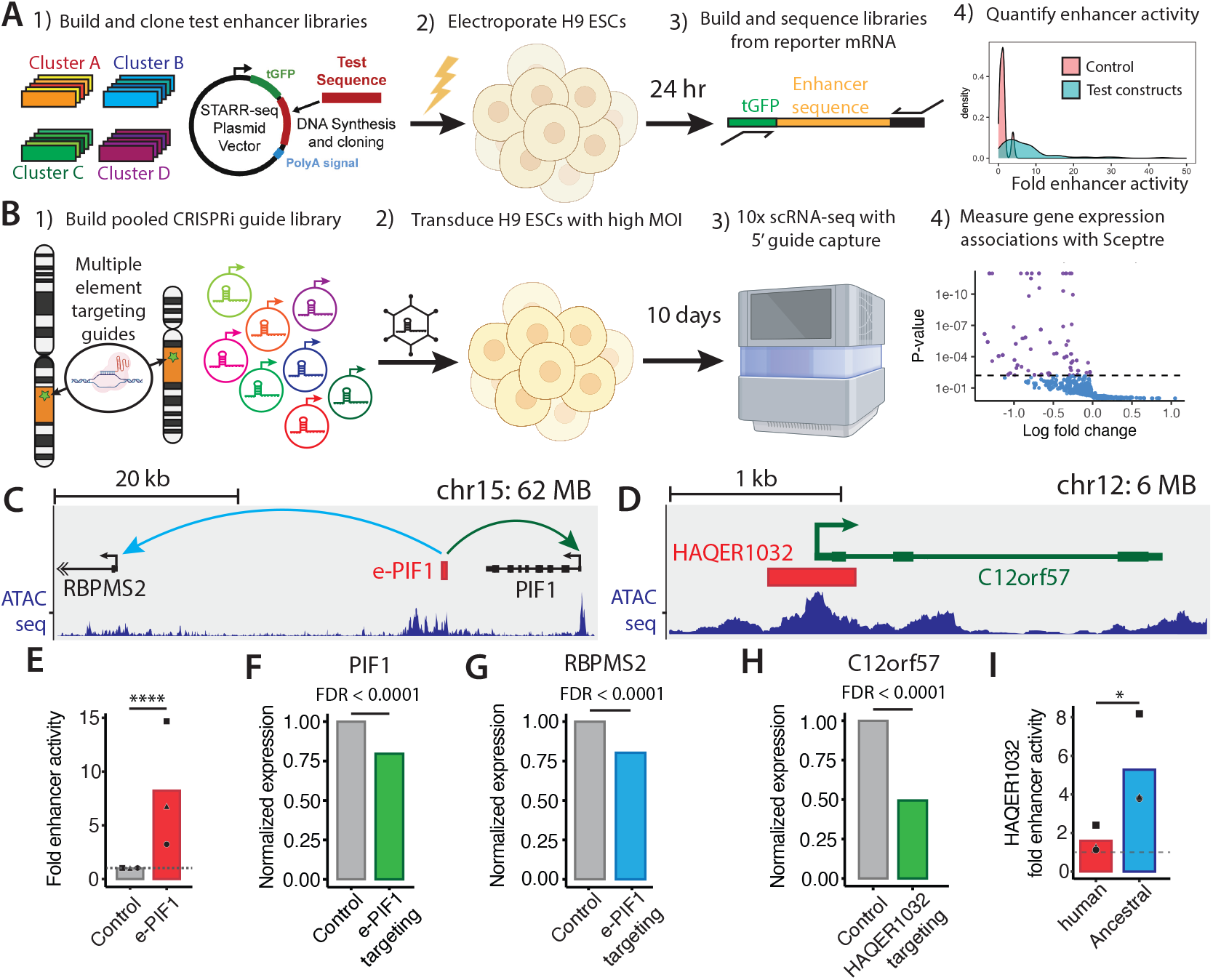
Orthogonal assays to identify enhancer activity. (**A**,**B)** Experimental design of STARR-seq and CRISPRi assays. We prioritized clusters for functional testing based on biochemical marks of enhancer activity and duplications being human-specific. For STARR-seq we performed three replicates. For CRISPRi we captured 20,000 cells. **(C)** Genomic context for the previously published enhancer, e-PIF1. e-PIF1 regulates *PIF1* and *RBPMS2*. **(D)** Genomic context of HAQER1032, which proximally regulates *C12orf57*. **(E)** Fold enhancer activity of e-PIF1 over negative controls from STARR-seq. Replicates are denoted by shape. (Stauffer’s method, ****: p *<* 0.0001) **(F**,**G)** Normalized *PIF1* and *RBPMS2* expression upon CRISPRi repression of e-PIF1 (Benjamini-Hochberg FDR < 0.1). **(H)** Normalized *C12orf57* expression upon CRISPRi repression of HAQER1032 (Benjamini-Hochberg FDR *<* 0.1). **(I)** Comparison of fold enhancer activity using STARR-seq between alleles of HAQER1032 representing the human-chimpanzee ancestor and modern humans. Replicates are denoted by shape. Dotted line is the average of negative controls (t-test; *: p *<* 0.05).

To complement the STARR-seq approach, we performed a high multiplicity of infection (MOI) CRISPR-interference (CRISPRi) screen to interrogate regulatory element activity in the endogenous context and identify the target gene(s) (Figure 2B). When feasible, we chose guide RNAs (gRNAs) that could target multiple paralogous elements within the same family, allowing us a greater ability to make comparisons between paralogous elements (see Methods). Historically, CRISPR gRNAs that target multiple locations in the genome were avoided because it was difficult to know with certainty which targeted location(s) were responsible for any observed transcriptional change. However, CRISPR regulatory element screening has become high throughput (Yao et al., 2024) and high MOI experiments that introduce multiple gRNAs into the same cell are becoming increasingly common (Gasperini et al., 2019). In the analysis of these screens, researchers only test for an association between a gene and a gRNA if the gene is within a particular genomic distance of the gRNA’s target site (Gasperini et al., 2019). This protects against false associations with the gene expression effects of other gRNAs within the same cell and reduces the multiple hypothesis testing burden. In our analysis, we tested for gene expression associations within 200kb on either side of each targeting gRNA location to focus on effects that are likely to be *cis-*acting direct targets of the noncoding elements (Figure S2B). A caveat of this analysis framework is that if two clustered elements from the same family are within 200kb of each other, we will not be able to infer which element(s) are responsible for any observed effect. In summary, we used a multi-targeting gRNA design, borrowing from analysis frameworks common to high MOI CRISPRi experiments, which allowed us to efficiently screen for the gene regulatory functions of duplicated noncoding elements at their endogenous loci.

Our CRISPRi experiment targeted a total of 107 clustered noncoding elements (from 24 clusters) with at least one gRNA (average of 3.13 guides per element). We then performed a *singleton analysis* were we tested for gRNA-gene expression associations at the level of individual gRNAs (Barry et al., 2024). We also performed a *union analysis* which tests for element-gene expression associations, pooling statistical power across all gRNAs targeting a noncoding element (see Methods). This increases the statistical power to detect smaller effects and/or effects in lowly expressed genes. Significant gRNA-gene associations were corrected to have a false discovery rate less than 0.1. Additionally, to ensure that we were controlling for false-positive associations, we randomly assigned non-targeting gRNAs to genes, using *sceptre* (Barry et al., 2021) calibration, and we detected no false positives in the singleton or union analyses (Figure S2C).

### STARR-seq and CRISPRi are complimentary

Along with negative controls, we also analyzed loci of known gene regulatory function in both the CRISPRi and STARR-seq assays to validate the complementary approach of the two assays. The first control was a distal enhancer regulating *PIF1*, e-PIF1, that was validated using CRISPR deletion (Barakat et al., 2018) (Figure 2C). In our STARR-seq assay e-PIF1 had greater than 8-fold stronger enhancer activity compared to negative controls (Figure 2E) (*p <* 0.00001). In our CRISPRi assay, we observed a 20% reduction in *PIF1* expression (L2FC = −0.33, raw p *<* 10^−38^, FDR *<* 0.0001) for cells with e-PIF1-targeting gRNAs (Figure 2F). This indicates that our CRISPRi assay can detect gene repression effect sizes that are relevant for developmental phenotypes (Richard et al., 2020; Muthuirulan et al., 2021). These results confirm that STARR-seq and CRISPRi screening can be combined to understand the regulatory activity of noncoding elements in isolation and at their endogenous location, as well as identifying their target gene(s).

Unexpectedly, we discovered that gRNAs targeting e-PIF1 also cause a 20% reduction in expression of *RBPMS2* (union, L2FC = −0.32, raw p *<* 10^−50^, FDR *<* 0.0001), a gene 35kb upstream of e-PIF1 (Figure 2G). This serves as a reminder that enhancers can regulate multiple target genes (Uyehara and Apostolou, 2023), and emphasizes that our CRISPRi screen is both able to detect these multiple associations and is unbiased in considering all nearby genes as potential targets.

A second control sequence was the promoter of *C12orf57*. This region has extensive human-chimpanzee divergence, and was identified as a *human ancestor quickly evolved region*: HAQER1032 (Figure 2D) (Mangan et al., 2022). CRISPRi gRNAs targeting HAQER1032, the promoter of *C12orf57*, caused a 59% reduction in *C12orf57* expression (union, L2FC = −1.27, raw p *<* 10^−50^, FDR *<* 0.0001) (Figure 2H).

Although HAQER1032 is annotated as a promoter, there is an opportunity to assay the distal regulatory ability of this DNA segment with STARR-seq. Previously, the extant human and ancestralized versions of this promoter element were tested with STARR-seq, with the ancestralized allele having significantly stronger enhancer activity than the extant human allele (Mangan et al., 2022). Our STARR-seq assay recapitulates this finding with the ancestralized allele having 6.81-fold stronger enhancer activity than the extant human allele (Figure 2I) (t-test, *p <* 0.024). This is consistent with proximal regulatory elements also having an ability to act distally in particular contexts (Andersson and Sandelin, 2020). Taken together, these results reemphasize that our STARR-seq and CRISPRi approaches are performing consistently with previously published results and can be used to understand both proximal and distal regulatory elements in both an isolated context and their endogenous context.

### Noncoding clusters are regulatory element families

Following this confirmation of our approaches, we next analyzed the activity of ESC clustered elements. First, we detected significant enhancer activity in 78 out of 112 clustered elements when tested in isolation with STARR-seq. When analyzing the CRISPRi data, we detected endogenous gene regulation at 28 of 107 clustered elements (Table S2). These 28 elements had a total of 38 significant regulatory connections, with 18 connections found in both singleton and union analysis strategies (Figure S2D). The different number of significant enhancers discovered between the two approaches is likely in part due to the different sensitivities of the assays, but may also reflect that some sequences have enhancer activity when tested outside of their chromatin context but do not actively regulate a gene at their endogenous location. Therefore, the tandem approach of STARR-seq and CRISPRi allows us to understand not only clustered elements with endogenous gene regulatory activity, but allows us to identify orphaned or chromatin masked clustered elements as well. Using these approaches, we validate many elements within noncoding clusters to function as gene regulatory elements and we therefore refer to validated clusters as regulatory element families (REFs). We proceed to explore case studies of REFs that highlight both the maintenance of regulatory relationships after duplication and multiple examples of genomic context-dependent regulatory activity.

### Maintenance of regulatory relationships

Many promoters maintain their regulatory activity following segmental duplication (Fraimovitch and Hagai, 2023). Maintenance of promoter activity following duplication can be grouped into two general categories with respect to the size of the duplication. The entire gene structure can be duplicated (i.e. promoter and all protein-coding exons) creating a duplicate copy of the original gene (Figure 3A). Alternatively, promoters can undergo smaller duplications where they are duplicated by themselves, or with only some of their accompanying exons, to become an alternative promoter of the same gene (Figure 3B), or a new gene based on a truncated version of the original gene (Figure 3C). We chose to focus on REF1 (Cluster 634) because it demonstrates both categories of promoter evolution.

**Fig. 3.**
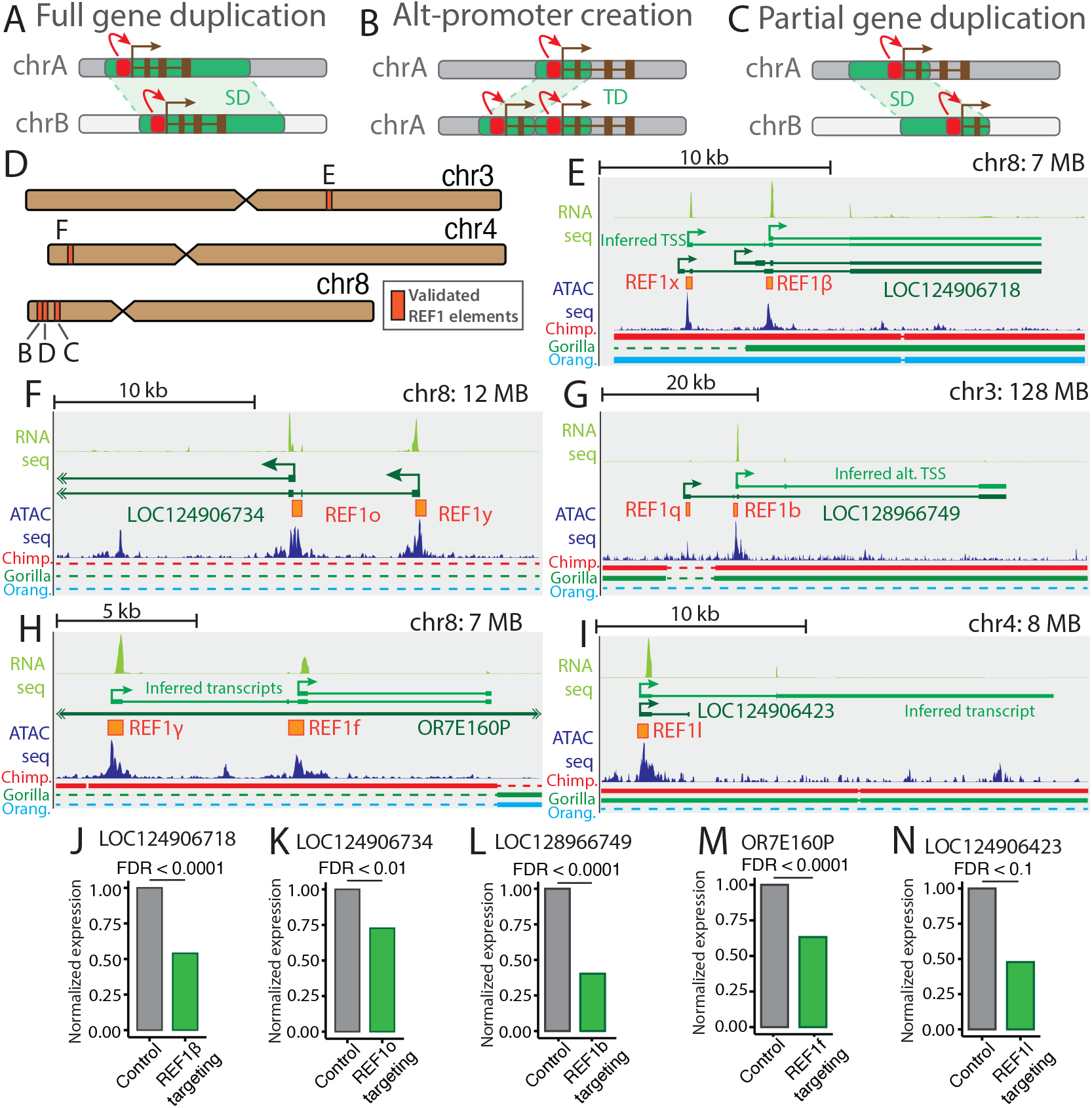
REF1 proximally regulates genes. **(A-C)** Models of complete or partial gene duplication. **(D)** Schematic of validated REF1 promoters, where letters correspond to figure panels showing additional genomic context. **(E-I)** Genomic contexts of REF1 promoter elements with significant CRISPRi signal. We used our 5’-capture RNA-seq from ESCs to refine the gene models near REF1 promoter genes, particularly with respect to transcription start site. To improve visualization, we show the inferred transcripts based on the ESC RNA-seq alignments, as well as the ESC RNA-seq alignments. Sequence alignments to T2T primate genomes are shown, with horizontal bars showing homology, and dashed lines showing alignment gaps (Chimp. = Chimpanzee; Orang. = Orangutan) **(J-N)** Normalized gene expression upon CRISPRi repression of REF1 elements (Benjamini-Hochberg FDR *<* 0.1).

The most straightforward case is when the entire gene structure is duplicated along with the promoter, likely resulting in a situation that is analogous to an upregulation of the original gene. REF1 is a family of 29 open elements where many are annotated as being promoters in the T2T gene set. While members of this REF are found on seven different chromosomes, 15 paralogs are located in the REPD and REPP SD clusters, which are 3.76 MB apart on chromosome 8 (Bosch et al., 2007) (Figure S3A). These SD clusters contain multiple gene families, including a *translation initiation factor IF-2-like* family. An example of a gene in this family is *LOC124906718* (Figure 3E), which underwent duplication via SD to create a human-specific copy, *LOC124906734* (Figure 3F) (Falker-Gieske, 2023). To verify that the promoters of these genes (which are members of REF1) are active, we targeted both with gRNAs in our CRISPRi screen. In cells that had gRNAs targeting REF1*β* (we use an additional letter to uniquely identify a member of a family), the downstream promoter of *LOC124906718*, we observed a 46% reduction in *LOC124906718* expression (Figure 3J) (L2FC = −0.89, raw p *<* 10^−5^, BH FDR < 0.0001). Similarly, cell with gRNAs targeting REF1o, the downstream promoter of *LOC124906734*, had a 27% reduction in *LOC124906734* expression (Figure 3K) (L2FC = −0.46, raw p *<* 10^−4^, BH FDR *<* 0.01). This duplication likely resulted in humans having greater expression of proteins with homology to *translation initiation factor IF-2-like* and this could potentially have effects on human-specific phenotypes since structural variation affecting these clusters cause developmental delay (Barber et al., 2008; Yu et al., 2010). While there are a number of similar instances in the literature of promoters maintaining activity upon full gene duplication (Zhang et al., 2022; Soto et al., 2025), this confirms that we are able to detect such events with our CRISPRi screen in ESCs.

Along with promoters being duplicated with their entire gene structure, they can also be duplicated in smaller tandem events where they continue to act on the original gene as an alternative promoter, or create a new gene based on a truncation of the original gene structure. Interestingly, the *translation initiation factor IF-2-like* genes from the first example not only have one REF1 element acting as a promoter, but have two REF1 elements, each acting as alternative promoters. Both overlap regions of open chromatin and serve as transcription start sites, based on 5’ RNA-seq alignments (Figure 3E,F). These alternative promoters are the result of an ancient tandem duplication event that formed a common duplicon and was subsequently spread around the genome through further SDs. While we are uncertain of the original tandem duplication event, we observed a human-specific tandem duplication on chromosome 3, creating the dual promoter configuration for a gene in humans, while the other great apes appear to have a single promoter driving expression of *LOC128966749* (Figure 3G). Using CRISPRi, we targeted REF1b, the downstream promoter of *LOC128966749*, and observed a 60% reduction in *LOC128966749* expression (Figure 3L) (L2FC = −1.31, raw p *<* 10^−20^, BH FDR *<* 0.0001). These examples show how tandem duplications of promoters can create novel isoforms and SDs of entire genes can expand gene families throughout the genome.

Along with tandem duplications creating alternative promoters, partial duplications can be inserted far from their source copy. Partial gene duplications have been shown to remain functional, and even create fusion transcripts (Dougherty et al., 2018). Therefore, we validated the REF1 promoter elements in two truncated *translation initiation factor IF-2-like* duplications that have created noncoding RNAs. *OR7E160P* is a gene duplication of the first two exons of the previously targeted *translation initiation factor IF-2-like* protein-coding genes, located at one end of the REPD cluster (Figure 3H). We used 5’-capture RNA-seq data to refine the gene annotation at the locus and predict that *OR7E160P* has the common dual-promoter structure previously observed with other REF1 genes (Figure 3H). When the downstream element in the tandem pair is targeted, REF1f, we observed a 37% reduction in OR7E160P expression (Figure 3M) (L2FC = −0.66, raw p *<* 10^−6^, BH FDR *<* 0.0001). There is an additional noncoding RNA gene with a REF1 promoter outside the REPP and REPD SD clusters, *LOC124906423*, within an SD on chromosome 4 (Figure 3I). Similar to other genes with REF1 promoters, *LOC124906423* expression was repressed by 52% in cells with promoter targeting gRNAs (Figure 3N) (L2FC = −1.07, raw p. = 0.0014, BH FDR *<* 0.1). We therefore suggest that REF1 elements have remained functional promoters even upon ancient or partial gene-duplication events. While there are number of similar examples of partially duplicated genes maintaining promoter activity following a segmental duplication (Fiddes et al., 2018b), the repression of five genes with REF1 promoters in response to a common CRISPRi gRNA stimulus shows the maintenance of promoter activity in ESCs after duplication (Figure 3D), and indicates that our multi-loci gRNA targeting framework can concurrently assess the *in vitro* function of multiple sites in the genome.

### Flexibility of functional element classes

In addition to detecting promoters maintaining their function as a proximal regulatory element, we also collected data on their ability to act more broadly as regulatory elements. For REF1 elements, we observed many maintaining their promoter activity upon duplication (Figure 3), but we also observed that all family members had significant enhancer activity when tested in isolation using the STARR-seq assay (Figure 4A). This led us to hypothesize that REF1 promoters may also be capable of distally regulating genes. To test this hypothesis, we investigated if gRNAs targeting REF1 promoters down regulated distally located genes and observed REF1 elements distally regulating genes at two loci. To begin understanding what mechanisms were controlling the gain of distal activity in some REF1 elements, we compared the STARR-seq activity of REF1 elements with distal activity from CRISPRi, to those REF1 elements tested with CRISPRi, but only showed proximal activity. We found that these two subsets of REF1 elements did not have significantly different enhancer activity from each other, as measured by STARR-seq (Figure S4A) (Wilcoxon, p = 0.59), suggesting that those promoters without endogenous distal enhancer activity may already be primed to perform distal regulation in a different genomic context.

**Fig. 4.**
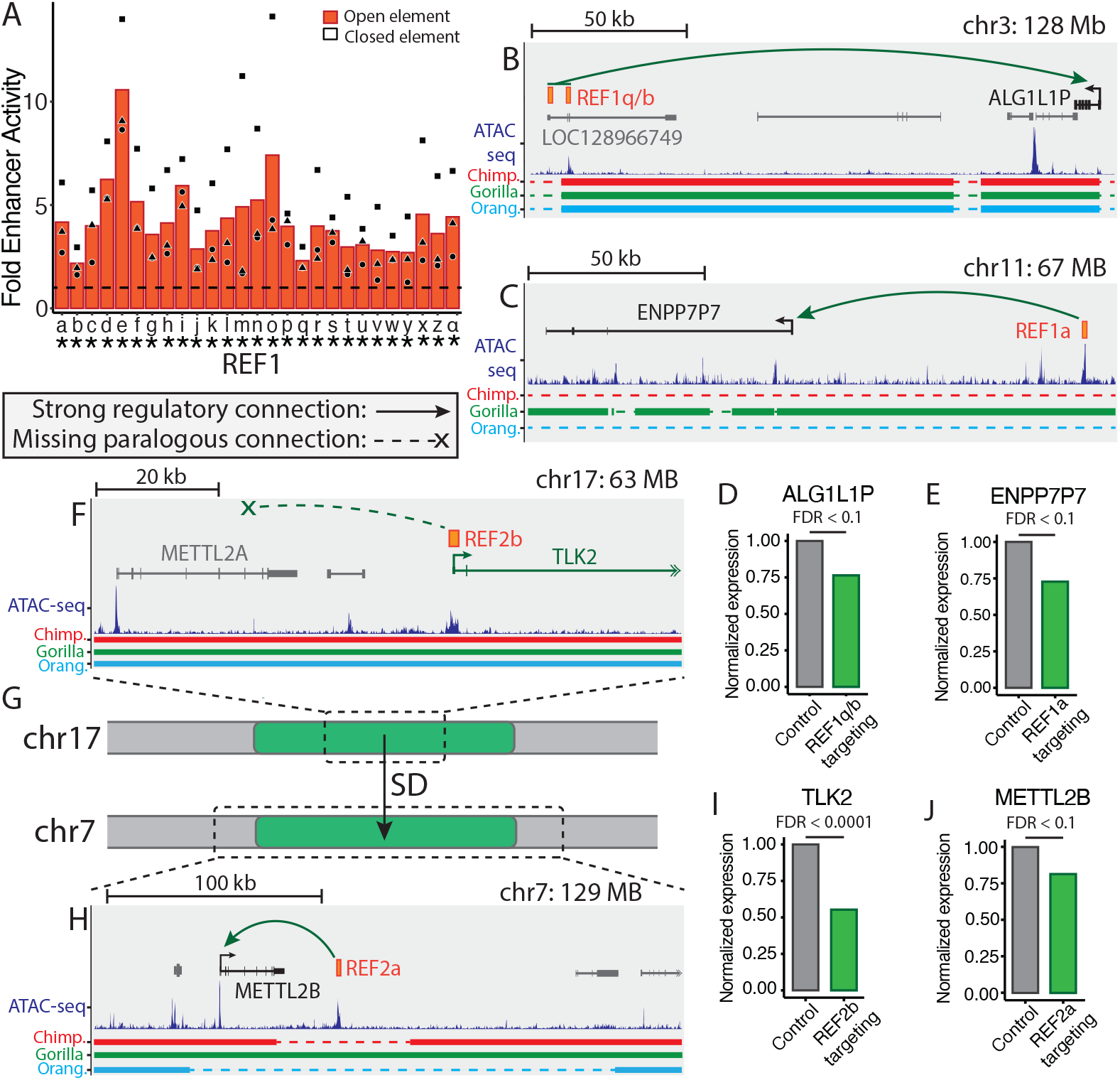
Proximal-to-distal element transitions after duplication. **(A)** Fold enhancer activity of REF1 elements over negative controls, measured by STARR-seq. Data point shape denotes replicate number. Dotted line is the average of negative controls (Stauffer’s method, *: p *<* 0.01) **(B**,**C)** Genomic contexts for REF1 elements with distal activity. Curved arrows represent distal regulatory interactions, which can be strong (solid line with arrow) or absent (dotted line with ex). Sequence alignments to T2T primate genomes are shown, with horizontal bars showing homology, and dashed lines showing alignment gaps (Chimp. = Chimpanzee; Orang. = Orangutan) **(D**,**E)** Normalized gene expression of *ALG1L1P* and *ENPP7P7* following CRISPRi repression of REF1q/b and REF1a, respectively (Benjamini-Hochberg FDR *<* 0.1). **(F)** Genomic contexts for the REF2b element. **(G)** Schematic showing the SD from chromosome 17 to chromosome 7. The dashed lines show then different viewpoint scales of the panels above and below. **(H)** Genomic contexts for the REF2a element. **(I**,**J)** Normalized gene expression of *TLK2* and *METTL2B* following CRISPRi repression of REF2b and REF2a, respectively (Benjamini-Hochberg FDR *<* 0.1).

First, when REF1q and REF1b, proximally-acting regulatory elements of *LOC128966749* (Figure 3E) are targeted with CRISPRi, we observed a 24% repression of a noncoding RNA 177kb downstream, *ALG1L1P* (Figure 4B,D) (L2FC = −0.39, raw p = 0.0016, BH FDR *<* 0.1). To asses the validity this long-range interaction where a promoter would also be acting as a distal regulatory element, we analyzed chromatin conformation data from ESCs (Krietenstein et al., 2020). We observed the REF1q and REF1b elements (promoters of *LOC128966749*) forming a stripe domain (Vian et al., 2018), which has an interaction with the promoter of the distally controlled gene, *ALG1L1P*, indicating that the promoters contact each other in 3-dimensional space (Figure S4B). Together, the REF1q/REF1b distal interaction provides evidence that promoters can also regulate distally located genes (Wei et al., 2022; Malfait et al., 2023; Wan et al., 2025).

Second, we further detected a REF1 element gaining distal activity upon duplication. The source of the duplication was a locus on chromosome 3, containing REF1k, which did not show distal activity upon CRISPRi repression. While there was not an annotated transcript beginning at this source locus, we detected 5-prime RNA-seq alignments mapping to REF1k, which supports its ability to act as a promoter (Figure S4C). Upon duplication to chromosome 11, REF1a overlaps a transcription start site and is located 82kb upstream of a noncoding RNA *ENPP7P7* (Figure 4C). In cells with REF1a targeting-guides, we observed a 27% reduction in *ENPP7P7* expression (Figure 4E) (L2FC = −0.46, raw p. = 0.0026, BH FDR *<* 0.1). These results indicate that this REF1 element, when duplicated, now acts as a distal regulatory element to regulate *ENPP7P7*. While the REF1 element on chromosome 11 was also not annotated as a promoter, we again analyzed 5-prime RNA-seq data from human ESCs and identified a small collection of reads that map to REF1a on chromosome 11. We used this data along with a gene model lifted from hg38 (Fiddes et al., 2018a) to add a gene model at this location (Figure S4E). When we included this gene model, named *LOC422-chr11*, in our reference gene set for the CRISPRi analysis, we detected very little expression and no significant downregulation when gRNAs targeted REF1a (L2FC = −0.28, raw p = 0.11, BH FDR *>* 0.1). This is likely due to the gene model only having a small number of reads that map to it, and therefore a lack of statistical power to detect down regulation. We propose that this example is not a complete transition from a promoter to enhancer upon duplication, but may represent us “catching evolution in the act,” during an element class transition, where enhancer activity has been gained, but promoter activity has not yet been fully lost.

We identified an additional example of a promoter transitioning to an enhancer in a different family, REF2 (Cluster 3641), with only two elements. The ancestral location, REF2b, is on chromosome 17 and is the promoter of *TLK2* (Figure 4F). When REF2b is targeted with CRISPRi, we observed a 45% reduction in *TLK2* expression, verifying its promoter activity (Figure 4I) (L2FC = −0.85, raw p. *<* 10^−5^, BH FDR *<* 0.0001). REF2b, and the surrounding 200kb, was segmentally duplicated to chromosome 7 (Figure 4G) at the time of the human-gorilla ancestor. This duplication created REF2a (Figure 4H). In the chimpanzee-bonobo ancestor, a subsequent 60kb deletion removed REF2a from the Pan lineage, leaving REF2a and its regulatory functions specific to gorillas and humans (Figure S4G). In humans, the duplication did not create an annotated *TLK2-like* gene copy on chromosome 7; however, we did detect distal regulatory activity at its new location. At the new location, REF2a distally regulates *METTL2B* (Figure 4H), a gene also in the original SD that resulted from the duplication of *METTL2A* and is involved in methylating tRNAs (Xu et al., 2017). When REF2a was targeted with CRISPRi, we observed an 18.5% reduction in *METTL2B* expression (Figure 4J) (L2FC = −0.30, raw p = 0.0049, BH FDR *<* 0.1). Surprisingly, the gRNA that effectively targeted the ancestral locus to repress *TLK2* did not significantly downregulate *METTL2A*. (Figure S4F) (L2FC = −0.13, raw p. = 0.18, BH FDR *>* 0.1), despite high expression of *METTL2A*. This suggests that the duplication of REF2b, the *TLK2* promoter, did not result in a *TLK2-like* protein-coding gene, but rather an enhancer for a nearby gene, *METTL2B*, within the SD. Similar to our analysis of the proximal-to-distal transition that occurred in REF1, we further investigated if a small amount of proximal activity could still be detected at REF2a. We investigated this possibility by adding a gene model, *TLK2-like-chr7*, at the paralogous location on chromosome 7 to the human T2T gene set that we used for the CRISPRi analysis (Figure S4H) (Methods). We also added a divergently-transcribed lncRNA, *LINC03072*, which was lifted from the hg38 gene set (Figure S4D). Both of these gene models are supported by ESC RNA-seq alignments (Figure S4H). When we re-ran the CRISPRi analysis, we observed a 39% reduction in *TLK2-like-chr7* expression when REF2a was targeted (Figure S4I) (L2FC = −0.71, raw p = 10^−12^, BH FDR *<* 0.0001) and also observed a 44% reduction in *LINC03072* expression (Figure S4I) (L2FC = −0.84, raw p = 10^−7^, BH FDR *<* 0.0001). These results indicate that while a functional protein-coding *TLK* gene copy was not maintained in the chromosome 7 SD, REF2a still has promoter activity, although *TLK2-like-chr7* has over four-fold lower expression than *TLK2* (Figure S4I). In summary, this is a second example of a promoter, without noticeable distal regulatory activity, that upon duplication gains distal activity and currently has only minimal activity as a promoter. Taken together, the results highlight not only examples of segmental duplications leading to gene regulatory rewiring, but propose that gene regulatory elements can readily transition between acting as promoters and enhancers, based on the genomic context in which they are placed.

### Duplicated enhancers rewire and facilitate regulatory innovation

After focusing on the gene regulatory consequences of promoter duplications, we proceeded to investigate the fate of distal elements after duplication. From our clustering analysis, we observed that REF3 (Cluster 867) is a family of 18 distal elements (Figure 5A), which we tested with STARR-seq and CRISPRi. We first characterized the enhancer activity of REF3 elements in isolation with STARR-seq. We observed that enhancer activity is generally conserved after duplication, with 14 of 18 elements having significant enhancer activity in STARR-seq (Figure S5A). While the elements with open chromatin in ESCs tend to cluster together by sequence similarity (Figure 5A), chromatin status is not significantly correlated with STARR-seq activity in REF3; however, there is a non-significant trend for open elements to show more activity (Figure S5B) (Wilcoxon, p = 0.083). This is consistent with both specific sequence features and open chromatin being required for endogenous enhancer activity (Peng et al., 2020; Sahu et al., 2022). These results are consistent with the genome containing a number of chromatin masked enhancers that already have the innate ability to act as distal regulatory elements if a further genomic change alters their larger context in a way that is more permissive of endogenous enhancer activity.

**Fig. 5.**
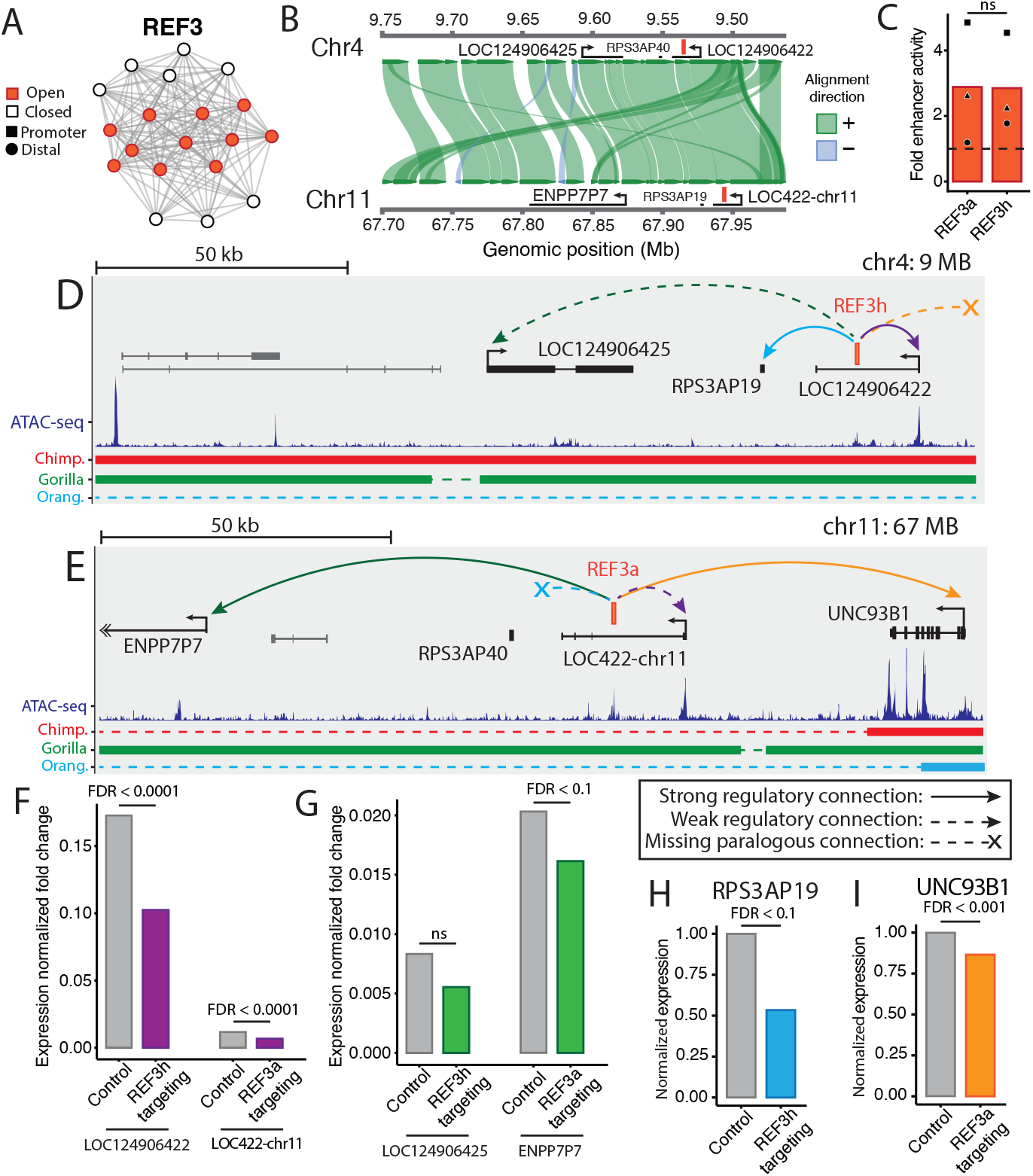
Distal regulatory rewiring in REF3. **(A)** REF3 visualized with edge lengths corresponding to percent divergence between noncoding element nodes. **(B)** Comparison of fold-enhancer activity of REF3a and REF3h, measured by STARR-seq. Dotted line is the average of negative controls (t-test, ns: p *>* 0.05). **(C)** Alignment of the chromosome 4 and chromosome 11 SDs, containing REF3h and REF3a, respectively **(D**,**E)** Genomic contexts for REF3a and REF3h. Curved arrows represent distal interactions, arrow color corresponds with CRISPRi fold-change plots below. Strong interactions are shown with a solid line, while weak interactions use a dotted line. Missing interactions are shown with a dotted line and an ex. Sequence alignments to T2T primate genomes are shown, with horizontal bars showing homology, and dashed lines showing alignment gaps (Chimp. = Chimpanzee; Orang. = Orangutan) **(F**,**G)** Fold-change gene expression plots showing conserved regulatory interactions when REF3 elements are repressed with CRISPRi in the paralogous SDs. Control and targeting gene expression bars are scaled by gene expression in all cells that did not receive a targeting gRNA, as calculated by Seurat (Hao et al., 2021) (Benjamini-Hochberg FDR *<* 0.1). **(H**,**I)** Normalized gene expression of *RPS3AP19* and *UNC93B1* when REF3h and REF3a are repressed with CRISPRi (Benjamini-Hochberg FDR *<* 0.1).

REF3 elements have been duplicated as part of larger SDs, which often included neighboring genes. Therefore, the REF3-containing SDs provide “natural experiments” by which to investigate how regulatory interactions between paralogous enhancer-gene pairs may be similar or different at various genomic loci. The family of SDs in which REF3 is contained was greatly expanded in copy number in the human-gorilla ancestor. 10 out of 18 REF3 elements are shared with gorilla but not with more distantly-related primates. Similar to REF1, we hypothesize that the REPD cluster is an ancestral location responsible for the source of REF3 duplications in the human-gorilla ancestor. This is because REF3 elements in the REPD cluster have synteny to primates more distantly related than gorilla, such as orangutan. When testing REF3 elements with CRISPRi, we detected enhancer activity at an ancestral location in the REPD cluster using both STARR-seq and CRISPRi, consistent with the ancestral state of REF3 elements has endogenous enhancer function (Figure S5C,D). We then focused further experiments on two REF3 elements within largely paralogous SDs on chromosome 4 and chromosome 11, responsible for creating REF3h and REF3a (Figure 5B). Both SDs occurred in the human-gorilla ancestor, however, the SD on chromosome 11 containing REF3a was potentially subject to incomplete lineage sorting, leading it to be shared with gorilla, but not chimpanzee. REF3h and REF3a have greater than 93% sequence identify and nearly identical enhancer strengths when measured in isolation with STARR-seq (Figure 5C) (t-test, p = 0.98). Therefore, these two SDs provide natural experiments to focus on if genomic context, rather than sequence-dependent enhancer strength, influences regulatory rewiring and interaction strength. The duplicated REF3 elements and paralogous genes may have conserved regulatory interactions at both paralogous locations or they may rewire and forge new regulatory interactions after duplication, both with genes inside the SD or genes already present in the locus.

There are multiple distal regulatory connections between REF3 elements and genes within their respective SDs, so comparing and contrasting these connections between the paralogous SDs on chromosome 4 (REF3h) and chromosome 11 (REF3a) can potentially identify examples of how regulatory interactions can change with context changes and evolutionary time. We found that while the interactions were generally conserved across paralogous locations, the strength of the regulatory interactions changed. The first example is a regulatory interaction in the chromosome 4 SD where REF3h is located within the first intron of a noncoding RNA, *LOC124906422*, which it regulates (Figure 5D). When REF3h is targeted with gRNAs, we observed a 44% downregulation of *LOC124906422* (L2FC = −0.83, raw p *<* 10^−14^, BH FDR *<* 0.0001) (Figure 5F, left). We originally believed that this distal regulatory connection was not conserved at REF3a in the chromosome 11 SD, since no homologous *LOC124906422-like* transcript existed within the human T2T gene set. However, when previously analyzing this locus (Figure S4E), we observed a small number of ESC RNA-seq alignments that supported the existence of a *LOC124906422-like* transcript, *LOC422-chr11* (Figure S5E). This led us to wonder if the regulatory interaction was maintained at a low level of expression and therefore we re-ran the CRISPRi analysis with *LOC422-chr11* included in the analysis. When REF3a was targeted, we observed a 43% reduction in *LOC422-chr11* (Figure 5E), indicating that the REF3-*LOC124906422-like* regulatory interaction is mirrored at the paralogous SDs (L2FC = −0.80, raw p *<* 10^−5^, BH *<* 0.0001) (Figure 5F, right). Though we observed a similar fold-change in *LOC124906422-like* gene expression upon repression of REF3 elements, the absolute magnitude of gene expression change upon repression is 14x higher in the REF3h-*LOC124906422* regulatory interaction (Figure 5F). We therefore propose a model where regulatory interactions between distal elements and paralogous genes may be conserved after SD, but regulatory connections can increase or decrease in strength after duplication, leading to differential expression of target genes.

Both of these REF3 elements, REF3a and REF3h, also regulate noncoding RNAs, within the SDs, 70kb away on their other side. We suggest that the enhancer-promoter interactions in these examples are conserved, due to the noncoding RNAs sharing a paralogous promoter sequence; however *ENPP7P7*, regulated by REF3a on chromosome 11 and *LOC124906425*, regulated by REF3h on chromosome 4, are transcribed on opposite strands (Figure 5D,E). In contrast to the previous example where REF3h had the stronger regulatory interaction, we observed REF3a having the stronger regulatory interaction with the noncoding RNA, *ENPP7P7*, on chromosome 11 (Figure 5E). This observation is consistent with the finding that REF3a and REF3h did not have significantly different enhancer activity when tested in isolation (Figure 5C), further suggesting that we are not observing inherently different enhancer activity or CRISPRi efficiency, but rather a degree of regulatory rewiring at the respective loci. When REF3a on chromosome 11 was targeted with CRISPRi, we observed a 20% downregulation in *ENPP7P7* expression (Figure 5G, right) (L2FC = −0.33, p = 0.0011, BH FDR *<* 0.1). Interestingly, *ENPP7P7* is also regulated by REF1a (Figure 4C,E), illustrating how large SDs can expand multiple regulatory element families with a single event, leading to multiple complex rewiring events at the new locus. In contrast to the REF3a-*ENPP7P7* regulatory interaction, we did not detect a significant interaction between REF3h and *LOC124906425* (Figure 5D) (L2FC = −0.59, raw p = 0.01, BH FDR *>* 0.1). However, the REF3h- *LOC124906425* interaction is trending towards significance, leading us to hypothesize that this is a conserved regulatory interaction albeit with an absolute change that is 1.5-fold weaker on chromosome 4 (Figure 5G, left). These results further support the model that while distal enhancer-gene regulatory interactions may be conserved after SD, their strengths of interaction are free to become stronger or weaker in a manner that is separable from the enhancer activity as measured in isolation by STARR-seq. Further, we hypothesize the observed weakening or strengthening of regulatory interactions after a recent duplication may reflect an intermediate evolutionary state where selection is acting to strengthen, weaken, or completely lose regulatory connections.

While we detected a number of conserved regulatory connections with differing strength between the two respective SDs, we also observed instances of regulatory rewiring due to the complete gain or loss of enhancer-gene connections. First, in the chromosome 4 SD, we observed a regulatory connection to a noncoding RNA, *RPS3AP19* (Figure 5D). When REF3h is targeted with CRISPRi, we detected a 46% downregulation in *RPS3AP19* (Figure 5H) (L2FC = −0.91, raw p. = 0.0019, BH FDR *<* 0.1). In contrast, the homologous transcript to *RPS3AP19, RPS3AP40*, was not expressed in ESCs, indicating that despite REF3a demonstrating enhancer activity, it does not contact the *RPS3AP-like* gene (Figure 5E). Therefore, we uncover complete regulatory rewiring between paralogous distal enhancers and genes within SDs, where regulatory interactions are completely gained/lost after duplication.

We additionally detected full regulatory rewiring at REF3a, generating a regulatory connection between REF3a and a protein-coding gene, *UNC93B1* (Figure 5E). Interestingly, *UNC93B1*, 60kb upstream of REF3a, was not duplicated along with REF3a but rather was already present in the locus on chromosome 11 pre-duplication. When REF3a is targeted with CRISPRi, we detected a 13% reduction in *UNC93B1* expression (Figure 5I) (L2FC = −0.21, p *<* 10^−5^, BH FDR *<* 0.001), indicating that regulatory rewiring after duplication is not restricted to genes within SDs but can rewire to regulate genes outside the SD. Consistent with human *UNC93B1* having a regulatory connection to a transcriptional enhancer not shared with chimpanzees, *UNC93B1* was identified as upregulated in human-chimpanzee allotetraploid iPSCs due to *cis-*regulatory differences (Figure S5F) (Gokhman et al., 2021). In total, we have identified five transcripts co-regulated by REF3 elements at three out of four CRISPRi-tested loci. Further, the majority of REF3 elements having STARR-seq activity suggests that REF3 may be controlling an even larger co-regulatory network than we were able to directly test in the CRISPRi screen. Of the loci we did test with CRISPRi, we have identified both the strengthening and weakening of existing connections, as well as the complete gain and loss of connections. This complete regulatory rewiring of enhancers in SDs involves both genes they were duplicated alongside within the SD and genes already present in locus before duplication.

### Segmental duplication hotspots expand functional regulatory element families

Since SDs are a major mechanism for creating and expanding noncoding element clusters (Figure 1G), places in the genome that are SD hotspots likely make outsized contributions to creating and expanding regulatory element families. Regions near the ends of chromosomes, referred to as subtelomeres, are prone to SDs and other structural variation (Mefford and Trask, 2002; Linardopoulou et al., 2005; Vollger et al., 2022). Consistent with these observations, we found that both open and closed ESC clustered element sets are enriched near the ends of chromosomes compared to ESC ATAC-seq peaks not in clusters (Figure 6A) (p *<* 10^−16^) (see Methods). Additionally, closed clustered elements are enriched near chromosome ends over open clustered elements (Figure 6A) (p *<* 10^−16^). This may be due in part to the telomere position effect (Baur et al., 2001), supporting that subtelomeric SDs contain a high proportion of “chromatin masked” regulatory elements (Peng et al., 2020; Sahu et al., 2022). Subtelomeric SDs commonly duplicate between subtelomeres, but also occasionally duplicate into euchromatic regions in the middle of chromosomes. We therefore hypothesize that subtelomeres expand families of functional elements, often in a chromatin masked state, and also facilitate the duplication of these family members into gene-rich euchromatic regions, where their regulatory potential is revealed.

**Fig. 6.**
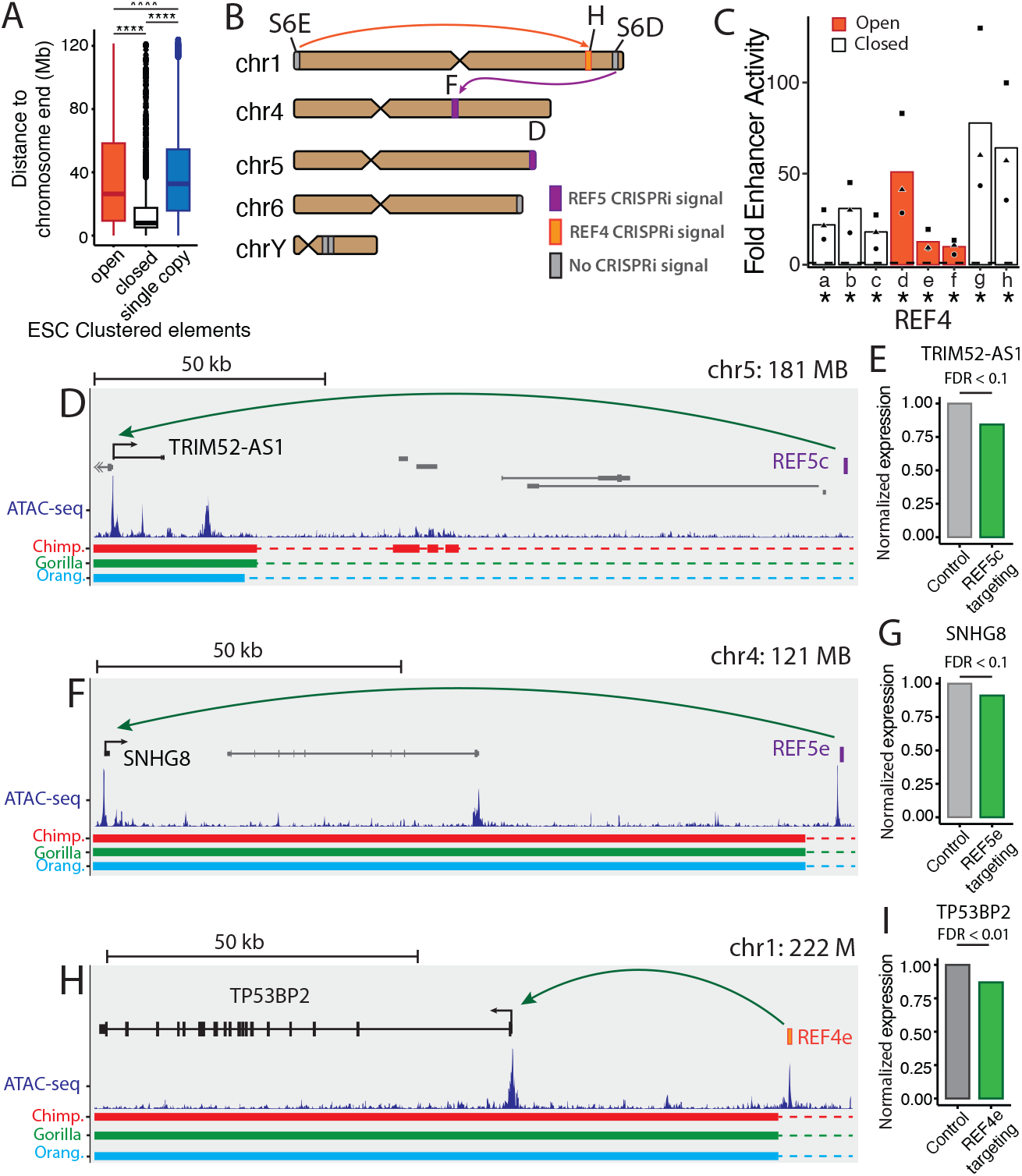
SD hotspots expand regulatory element families. **(A)** Distance of ESC open clustered elements, closed clustered elements, and single copy elements to chromosome ends in the human T2T reference assembly (Wilcoxon, ****: p *<* 0.0001). **(B)** Schematic of human chromosomes with REF4 and REF5 elements. Colors denote loci with CRISPRi-responsive elements, arrows reflect the source of duplication, and letters correspond to figure panels showing additional genomic context. **(C)** Fold enhancer activity of REF4 elements over negative controls, measured by STARR-seq. Data point shape denotes replicate number. Dotted line is the average of negative controls (Stauffer’s method, *: p *<* 0.01) **(D**,**F**,**H)** Specific genomic context of selected REF4 and REF5 elements. Curved arrow represents a distal gene regulatory interaction. While there was not sufficient ATAC-seq read-depth to call a significant open chromatin peak at REF5c, panel D, we observed a nonsignificant read pile-up, suggesting that this loci may be weakly open in ESCs. Sequence alignments to T2T primate genomes are shown, with horizontal bars showing homology, and dashed lines showing alignment gaps (Chimp. = Chimpanzee; Orang. = Orangutan). **(E**,**G**,**I)** Normalized *TRIM52-AS1, TP53BP2, SNHG8* expression when their respective distal enhancers are targeted with CRISPRi (Benjamini-Hochberg FDR *<* 0.1).

We observed two separate regulatory element families, REF4 and REF5, contained within the same segmental duplicon and commonly found within subtelomeres. Copy number of both REF4 and REF5 families have increased throughout primate evolution. The orangutan genome has only a single element for each of REF4 and REF5 at a primate-conserved locus on chromosome 1, located 6 Mb from the end of the chromosome (Figure S6A). In humans, REF4f and REF5d are at this ancestral locus and both have open chromatin signal and STARR-seq enhancer activity (Figure 6C, S6C, S6D), suggesting that the ancestral state of both families has gene regulatory potential. In the human-gorilla ancestor, this locus was duplicated into a subtelomeric region, where it underwent subsequent lineage-specific expansion in the gorilla, chimpanzee, and human genomes (Figure S6A). In the human genome, both REF4 and REF5 are comprised of eight elements each (Figure S6B). Four members of both REF4 and REF5 are located within 6 Mb of the ends of chromosomes and two members of each family are within an SD cluster on the q-arm of the Y chromosome (Figure 6B). This provides support for SD-prone regions as a mechanism for rapidly expanding regulatory element families and highlights the benefits of telomere-to-telomere genome assemblies.

We detected widespread enhancer activity of both REF4 and REF5 elements when tested in isolation with STARR-seq. Particularly, REF4 had the strongest STARR-seq enhancer activity of any tested cluster, with activity ranging from 9.9 to 77.8-fold stronger enhancer activity over controls (Figure 6C). We detected more modest enhancer activity in REF5: 1.4 to 2.6-fold stronger enhancer activity over control, with only two out of eight REF5 members having significant enhancer activity (Figure S6C). This result is consistent with regulatory elements maintaining the potential for enhancer activity in these SD-prone regions even though they can be chromatin masked. While we did not detect significant CRISPRi associations for most REF4 and REF5 elements in subtelomeres, we did detect a gene expression association between REF5c in a chromosome 5 subtelomere and an anti-sense noncoding RNA, *TRIM52-AS1*, indicating that elements in this family can have endogenous enhancer activity even in subtelomeres (Figure 6D,E) (L2FC = −0.25, raw p. = 0.0061, BH FDR *<* 0.1). These results suggest that while subtelomeric SDs have expanded elements with intrinsic enhancer activity, they are often in genomic contexts that are not permissive to endogenous gene regulation. Genomic regions with elevated mutation rates likely made outsized contributions to functional mutations on the human lineage (Xie et al., 2019; Mangan et al., 2022). Consistent with subtelomeric regions being prone to structural variation, we observed two human-specific SDs, both containing a REF4 and a REF5 element, that were duplicated into gene-rich, euchromatic regions (Figure 6F,H). While most elements in REF4 and REF5 are chromatin masked, we observed robust ESC ATAC-seq signal at these euchromatic elements. We therefore wondered if the open chromatin signal at these duplicated elements reflected a change from closed, chromatin-masked elements to active elements with endogenous regulatory activity. The human-specific SD on chromosome 4, which created REF5e, (Figure 6F) duplicated from the primate-conserved REF4f/REF5d locus on chromosome 1. While the ancestral REF5d element on chromosome 1 had open chromatin signal and was robustly targeted by gRNAs (Figure S6D), we did not detect any endogenous gene regulatory activity, suggesting that REF4f is an example of an orphaned enhancer. However, when REF5e within the human-specific SD on chromosome 4 is targeted with an active gRNA, we observed a 9% decrease in *SNHG8* expression, a small nucleolar RNA 145kb away outside of the human-specific SD. This result suggests that when REF5 elements are placed in the proper context they can gain endogenous regulatory activity (Figure 6G) (L2FC = −0.13, raw p. = 0.0016, BH FDR *<* 0.1).

Further, we observed a human-specific SD on the q-arm of chromosome 1 (Figure 6H) that duplicated from the subtelomeric region on the p-arm of chromosome 1 (Figure S6E). At the source copy in the p-arm subtelomere, REF4b does not have open chromatin or endogenous regulatory activity (Figure S6E), but when tested in isolation, has roughly 30-fold greater enhancer activity over negative controls (Figure 6C) (p *<* 0.00001), suggesting that this closed REF4 element is chromatin masked. However, when the euchromatic REF4e is targeted with an active gRNA, we observed a 13% reduction in *TP53BP2* expression, a gene 45kb upstream not contained within the SD (Figure 6I) (L2FC = −0.20, raw p = 0.00042, BH FDR *<* 0.1). Taken together, these results suggest that chromatin masked elements in subtelomeres can retain ancestral enhancer activity and undergo rapid lineage-specific expansion. Subsequently, we observed SD-prone subtelomeric loci deposit these previously chromatin-masked elements into euchromatic regions where their ability to endogenously regulate genes was “rescued” and contributed to human-specific regulatory rewiring at the locus.

Interestingly, both *TP53BP2* and *SNHG8* have established roles in cancer biology. *TP53BP2* is a tumor suppressor gene which promotes apoptosis in both *p53*-dependent and *p53*-independent pathways, and is downregulated in many malignant tumors (Huo et al., 2023). *SNHG8* plays roles in cell proliferation, migration, and in the epithelial-mesenchymal transition during development (Ghafouri-Fard et al., 2023; He et al., 2022). Consequently, *SNHG8*-dependent mechanisms are often co-opted in cancer and the upregulation of *SNHG8* expression in tumors promotes their proliferation and migration (Yuan et al., 2021; Ghafouri-Fard et al., 2023). The observed novel human connections to these genes highlight that human-specific SDs may have modulated our risk for diseases, such as cancer, by rewiring the regulation of disease-associated genes in ways that were beneficial or deleterious (Benton et al., 2021; Vaill et al., 2023).

### Regulatory element families across iCM development

Following the functional validation of SD-mediated noncoding clusters as regulatory element families in ESCs, we aimed to expand our understanding of how regulatory element families may function in other cell types. Since gene co-regulation is known to be especially important during development (Razin et al., 2021), we hypothesized that regulatory element families may contribute to gene co-expression modules in a model of cardiomyocyte (CM) differentiation. In addition to the H9 ESC ATAC-seq dataset, we further analyzed ATAC-seq datasets from cardiac mesoderm and cardiomyocyte (days 0, 4, and 30 of *in vitro* cardiomyocyte differentiation; see Methods) (Liu et al., 2017). Our clustering analysis on elements that have chromatin accessibility in at least one time point during (CM) differentiation identified 2100 families containing 7614 open chromatin peaks (Table S3). Of the 7614 total open elements, 3837 elements (50%) had open chromatin at only one time point, indicating a subset of elements in families likely act at only a specific set of stages during CM differentiation.

We identified families primarily open at a single time point to investigate how regulatory element families can contribute to co-regulatory networks that facilitate transcriptional switches during cardiomyocyte differentiation. This analysis yielded 1071 “time point-specific families” (TP-families), families in which more than half of open elements were open at a single time point during CM differentiation (Figure 7A). Similar to the previously identified ESC regulatory element families, SDs are a mechanistic force creating TP-families (Figure 7B) (fold enrichment = 6.24, p *<* 10^−20^). Since elements in families share intra-cluster DNA sequence similarity, we hypothesized that specific transcription factor binding motifs would be enriched in TP-family elements at a specific time point compared to other time points. Illustrating this, we detected enrichments of relevant transcription factor binding motifs at all tested timepoints (Figure 7C) (Table S4) (see Methods). Of particular interest, the transcription factor *ZEB1*, required for cardiac mesoderm differentiation into mature CMs (Liu et al., 2017; Ninfali et al., 2023), is 1.29-fold enriched in cardiac mesoderm TP-clustered elements (*p*_*adj*_ *<* 10^−50^). *FOS-JUN* motifs, two subunits of the *AP-1* transcription factor complex (Bejjani et al., 2019), crucial for post-natal cardiomyocyte maturation (Beisaw et al., 2020; Zhang et al., 2023), were similarly enriched in cardiomyocyte TP-families compared to other time points (Fold enrichment = 5.25, *p*_*adj*_ *<* 10^−50^). Taken together, we suggest that SDs create time point-specific regulatory element families that bind *trans-*acting factors expressed at that time point in order to switch on and off batteries of genes to influence cardiomyocyte differentiation.

**Fig. 7.**
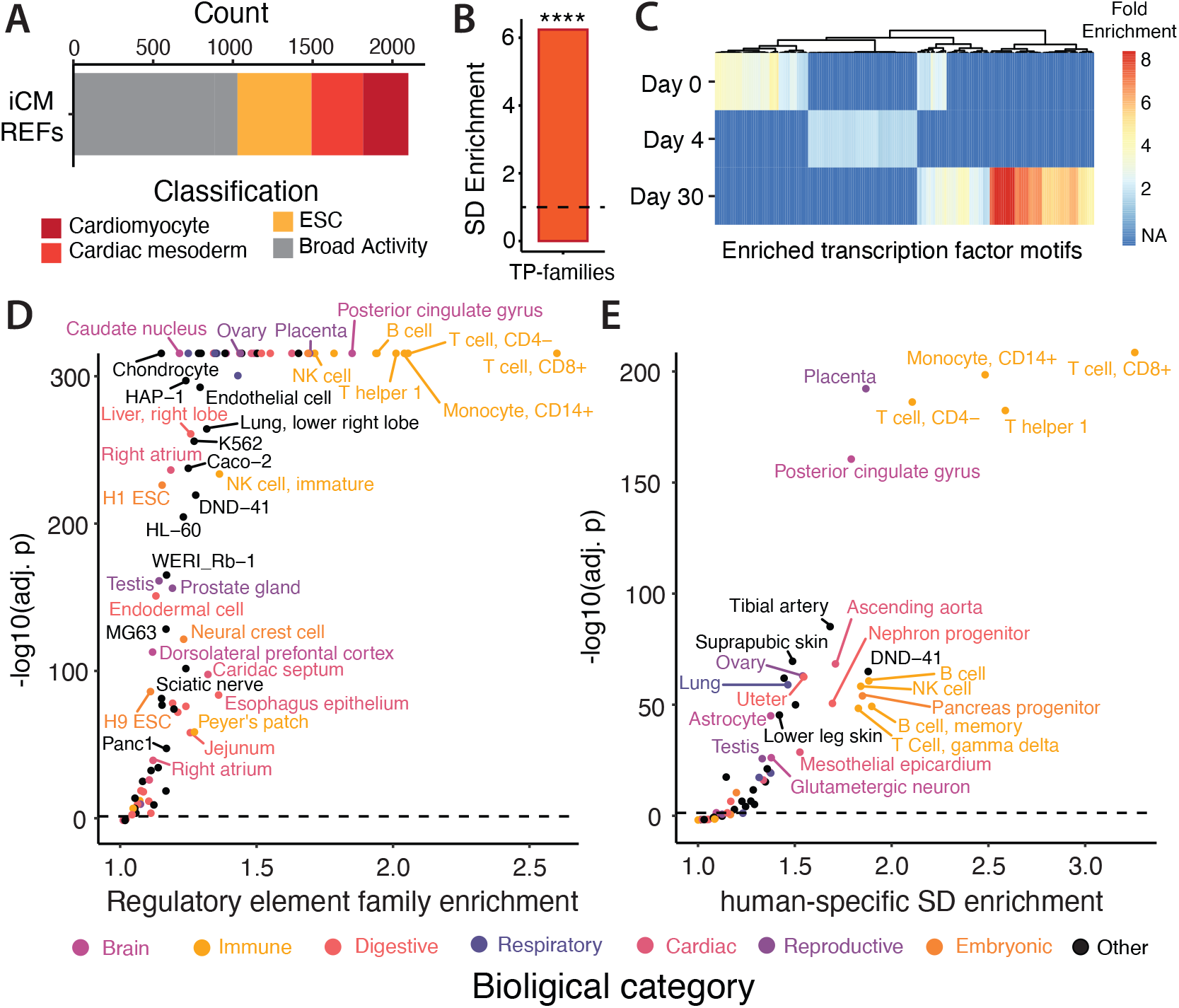
Biological roles of regulatory element families in diverse cell types. **(A)** 2100 total regulatory element families in *in vitro* cardiomyocyte development (iCM REFs), colored by active time point(s). **(B)** Enrichment of human T2T SDs for time point-specific family elements (TP-families) from iCM differentiation. (Binomial; ****: p *<* 0.0001). **(C)** Transcription factor binding motifs significantly enriched in TP-clusters at a certain timepoint compared to other timepoints. Timepoints of *in vitro* cardiomyocyte differentiation: ESC = day 0, Cardiac mesoderm = day 4, cardiomyocyte = day 30 (Binomial, Bonferroni-adjusted p *<* 0.05). **(D**,**E)** Enrichments of open chromatin regions from 154 ENCODE cell types in regions of human T2T self-paralogy or human-specific SD, respectively. We plotted data points with enrichments greater than one. For simplicity, samples that had a high degree of biological overlap with other tested samples (e.g. similar T cell states) were omitted from the plot, but not multiple hypothesis correction, prioritizing the sample with the higher enrichment. Data points are colored by biological meta-categories. Dashed line represents significance threshold (Binomial, Bonferroni-adjusted p *<* 0.05).

### Immune cell types and brain subregions are enriched for regulatory element families

Beyond cell types on the developmental lineage to cardiomyocytes, we were curious what cell types and tissues have been most affected by the creation of regulatory element families in recent evolution, including human-specific changes. To answer this question, we re-aligned chromatin accessibility data from 154 different ENCODE cell types to the T2T human genome (see Methods) (Table S6) (ENCODE Project Consortium, 2012). We observed regulatory element families, including human-specific element expansions, enriched in chromatin accessibility datasets from numerous immune-related cell types, including T-cells, monocytes, and B-cells (Figure 7D,E) (T-cell enrichment: REF 2.60 fold, *p*_*adj*_ *<* 10^−50^, human-specific SD 3.26 fold, *p*_*adj*_ *<* 10^−50^; monocyte enrichment: REF 2.04 fold, *p*_*adj*_ *<* 10^−50^, human-specific SD 2.48 fold, *p*_*adj*_ *<* 10^−50^; B-cell enrichment: REF 1.94 fold, *p*_*adj*_ *<* 10^−50^, human-specific SD 1.90 fold, *p*_*adj*_ *<* 10^−50^) (Table S5). This is consistent with the observation that genomic regions involved with immune function are associated with structural variation, leading to fixed differences between humans and chimpanzees (Anzai et al., 2003; Sequencing and Consortium, 2005) and polymorphic structural variants in the human population (Zhang et al., 2021; Logsdon et al., 2025). Clustered elements in immune cell types may play a role in the rapid induction of gene co-expression upon an immune challenge, similar to how many transposable element-mediated co-regulatory networks function (Chuong et al., 2016; Horton et al., 2023). These observed enrichments are consistent with the immune system rapidly evolving in humans as well as the immune response benefiting from gene co-regulation.

In addition to immune cell types, we also observed that open chromatin regions in the *posterior cingulate gyrus* are 1.85-fold enriched for being contained within regulatory element families (Figure 7D) (*p*_*adj*_ *<* 10^−20^), including a 1.79-fold enrichment for human-specific element expansions (Figure 7E) (*p*_*adj*_ *<* 10^−20^). This is consistent with the posterior cingulate gyrus having numerous human-specific gene regulatory changes, potentially influencing cell type functions and proportions compared to other primates (Caglayan et al., 2023). The posterior cingulate gyrus may be important for human-specific behaviors due to its proposed function in emotion, memory, and the sense of self (Guterstam et al., 2015; Finlayson-Short et al., 2020; Foster et al., 2023). Further, the posterior cingulate gyrus is thought to underpin susceptibility to neurological disorders such as schizophrenia and Alzheimer’s disease (Leech and Sharp, 2014), of which humans may be especially suseptible. Taken together, we suggest that gene co-regulation by families of SD-duplicated regulatory elements play roles in a diversity of biological processes, including human-specific changes to immune response and brain function.

## Discussion

In this manuscript, we provide supporting evidence for the importance of segmental duplications in shaping regulatory element families (REFs) in the human genome. Our functional analysis of REF elements with STARR-seq and CRISPRi uncovers diverse fates of regulatory elements after duplication. We observed conservation of existing regulatory relationships where duplicated regulatory elements either continue to act on the same gene as before, or act on the duplicated version of the original gene (Figure 8A). However, we also observe more complex changes upon regulatory element duplication. We observed promoters gaining distal activity while reducing their proximal activity, likely in the process of transitioning from a promoter to an enhancer upon duplication (Figure 8B). We also observed regulatory elements rewiring to form novel regulatory connections with a gene that it did not previously regulate. We observed this both in the context of active enhancers maintaining their function but changing the gene on which they act (Figure 8C), as well as “masked” or “orphaned” enhancers that had the inherent ability to regulate transcription, although without a target gene, gain a target gene and endogenous function upon duplication (Figure 8D). We hypothesize that the observed regulatory rewiring after segmental duplication is a mechanism by which transcripts can be brought into co-regulatory networks, as well as removed from them. The multiple examples of these changes on the human lineage provides evidence that regulatory rewiring after structural variation may be a common mechanism of gene regulatory evolution (Keough et al., 2023), and that in addition to specific sequence features, the genomic context of a regulatory elements is determinant of a regulatory element’s ultimate function.

**Fig. 8.**
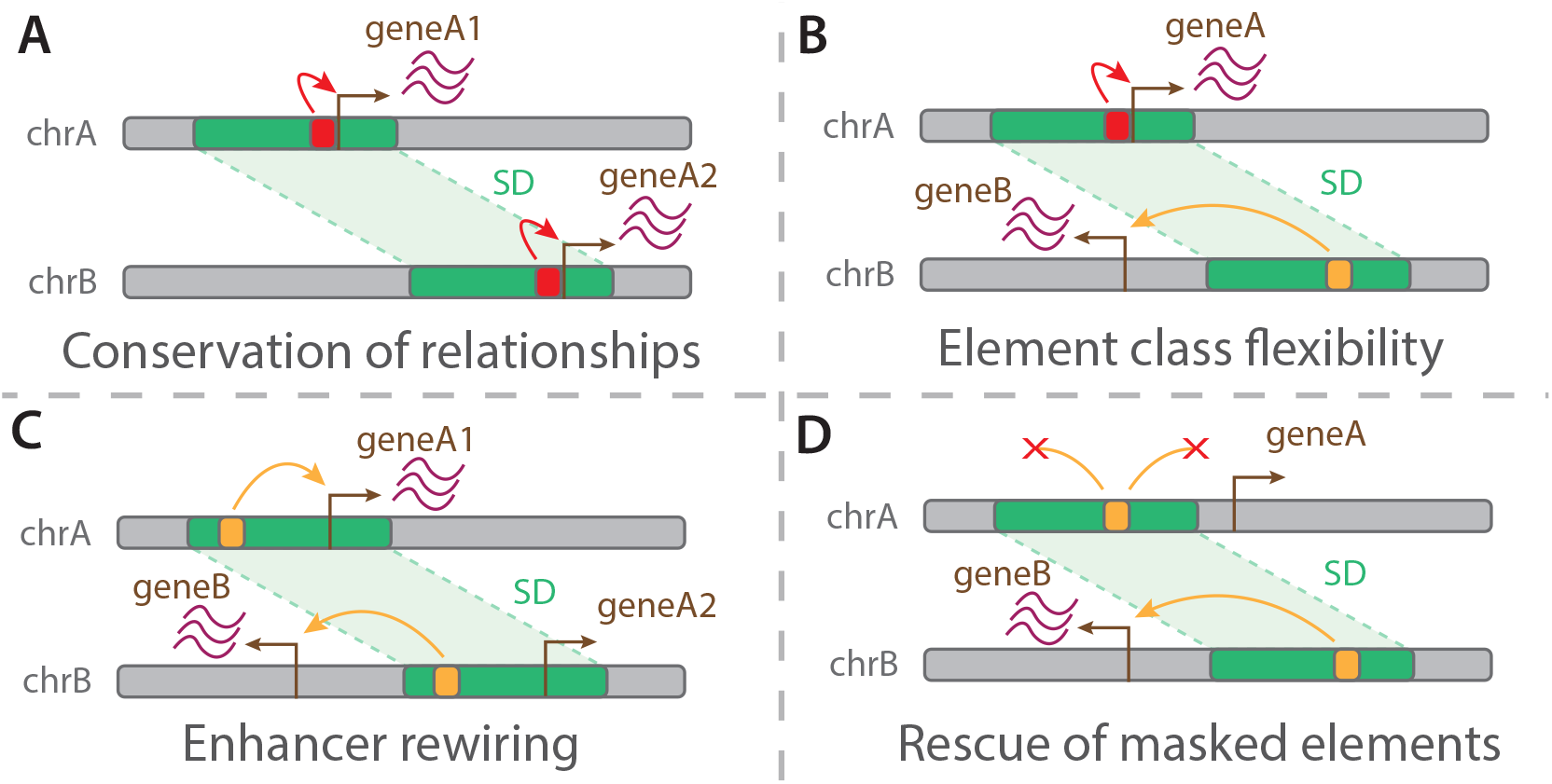
Observed fates of duplicated regulatory elements (A-D) A model showing four observed fates of regulatory elements after segmental duplication. Red elements signify proximally-acting elements, while yellow elements are distally-acting.

Similar to other recent works studying enhancer function with STARR-seq or MPRA-based assays, we observed masked enhancers that have activity when tested in isolation, but have no native open chromatin signal (Peng et al., 2020; Sahu et al., 2022). As has been previously suggested, there are multiple levels of chromatin regulation on top of DNA sequence features that matter for endogenous enhancer activity (Sahu et al., 2022). An example from this manuscript is elements within REF4 and REF5, which have strong STARR-seq activity, but have closed chromatin at their endogenous locations near the ends of chromosomes. We hypothesize that these subtelomeric enhancers are subject to the telomere positions effect (Baur et al., 2001) or other silencing mechanisms to repress their chromatin accessibility and therefore ability to act at enhancers at that locus. However, upon duplication to a gene-rich euchromatic locus, these REF4 and REF5 elements display open chromatin and actively regulate nearby genes, showing that duplication into a locus more permissive of endogenous gene regulation can rescue their function. These results also highlight how the dual approach of an episomal enhancer assay, such as STARR-seq, and an assay of endogenous enhancer activity, such as CRISPRi, can be combined to tell a richer story than either approach on it’s own.

In addition to masked enhancers, we also observed orphaned enhancers that were STARR-seq positive with open chromatin at their endogenous locus, but did not function as the regulatory elements of a gene based on our CRISPRi data. An example of this in our data is the primate-syntenic elements REF4f and REF5d, both elements which have open chromatin signal and were robustly targeted with CRISPRi gRNAs (Figure S6D). There are a few possibilities that would explain this observation. First, STARR-seq is potentially more sensitive than CRISPRi and is able to detect enhancer activity where the repression of the same enhancer with CRISPRi may cause only a very slight repression of a target gene or even is compensated for by other *cis-*acting elements (Hong et al., 2008). Additionally, due to differences in gRNA efficiency, it is possible that none of our selected gRNAs efficiently targeted the REF member, and therefore no repression occurred. However, a thorough case study in the literature observed duplications causing an endogenously active regulatory element to be orphaned (Franke et al., 2016). The recurrent duplication of a *SOX9* enhancer which serves to pathogenically upregulate *SOX9*, occasionally gets duplicated along with a topologically associating domain boundary. In these cases, a new topologically associating domain, including the duplicate enhancer, is created at the destination locus.

This serves to orphan the enhancer and prevents a pathogenic *SOX9* upregulation. Though in our data orphaned enhancers are rescued, not explicitly created as in the literature case study, we suggest that orphaned enhancers exist normally in the human genome, and the genomic context at the destination locus is a major determinant of endogenous regulatory activity and rewiring. A future analysis of STARR-seq positive and CRISPRi negative duplicated elements with their broader genomic context in mind, including 3D genome folding and the annotation of boundary elements, will be useful for providing additional evidence to this hypothesis.

An interesting finding from the functional study of REF elements is the apparent proximal-to-distal element transitions after duplication. We hypothesize that regulatory elements are primed for neofunctionalization after duplication, similar to protein coding genes (Ohno, 1970). We expect that this may materialize as elements switching the cell type in which they are active, changing the strength of regulation, or as we observed, switching between proximal and distal regulatory mechanisms. In the case of proximal-to-distal element transitions, we hypothesize that the intrinsic DNA-binding activity of promoter elements allow them to more easily gain distal activity compared to previously non-functional sequence. However, the mechanisms by which this occurs remains unknown. The effect of sequence changes on gene regulatory function has been the subject of considerable research through high-throughput reporter assays (Duan et al., 2023; Morova et al., 2023; Xiao et al., 2024); however, our work suggests that genomic context is an additional variable that warrants further study. REF elements that are at high copy number in genomes provide natural experiments to interrogate the effect of both genomic context and sequence features on regulatory function. A more comprehensive characterization of how REF elements endogenously regulate genes in each duplicated locus will not only yield a more complete understanding of regulatory element neofunctionalization, but may elucidate further rules of how enhancers find and regulate promoters of genes.

We cannot make a direct comparison between TE-derived and SD-derived regulatory element families due to the different programs by which the sets of families are ascertained. However, the overrepresentation of REF elements in immune-related cell types suggests some functional overlap between TE-derived and SD-derived regulatory element families (Chuong et al., 2016; Horton et al., 2023). It is also clear from existing work that TE-derived regulatory element families, on the order of tens of thousands of elements, are much larger on average than our SD-derived REFs (Feschotte, 2008), perhaps hinting at separate types of regulatory changes that are possible with the two evolutionary mechanisms. Our identified REFs are generally on the order of two to tens of elements, and therefore are likely to control smaller gene modules. Additionally, TE duplication likely duplicates discrete, single elements, while a single SD can duplicate many regulatory elements in a single event. We hypothesize that large SDs can lead to plietropic and large-scale rewiring events, as we observed with REF1 and REF3 elements often being contained within the same SDs.

We observed regions prone to structural variation, such as subtelomeres, creating recent human-specific duplications that harbor regulatory elements that underwent regulatory rewiring and had functional consequences at their destination loci (Figure 6). We expect that these SD-prone regions create human-polymorphic SDs that influences regulatory element variation within human populations. A limitation of our current analysis is that we were restricted to a small number of available human and primate telomere-to-telomere reference genomes. While we ensured that SDs in our human-specific SD dataset were fixed within three diverse human genomes, this is not comprehensive when considering the breadth of human variation. However, the number of high quality human reference genomes is growing, and human polymorphic SDs and structural variants have been well documented in these genomes (Vollger et al., 2022; Rocha et al., 2024; Logsdon et al., 2025). We hope that integrating regulatory element analyses into this growing body of work will shed light on functional consequences of human polymorphic structural variation related to health and disease.

A subset of regulatory element copy number variants, both between species and within the human population, are likely to contribute to disease. In our manuscript we highlight a human-specific regulatory connection between a duplicated enhancer and a noncoding RNA oncogene (Yuan et al., 2021; Ghafouri-Fard et al., 2023) and uncover human-specific open chromatin regions enriched in a brain subregion associated with schizophrenia susceptibility (Foster et al., 2023). It is likely that the large-scale regulatory rewriring after segmental duplication, along with gene copy number changes, makes these mutations especially pleiotropic. We propose that this pleiotropy has made segmental duplications likely to underlie disease, in part due to evolutionary conflicts where a single segmental duplication results in both advantageous and deleterious regulatory rewiring.

### Limitations

The human T2T reference genome was only recently published and it is still being actively annotated both at the level of gene models, and functional genomic data sets that were originally analyzed in the context of incomplete assemblies. We were analyzing genomic regions with a high degree of paralogy and that have only recently been resolved (Nurk et al., 2022), which means there has been less time to refine their annotations. Multiple times we observed 5-prime RNA-seq data mapping to regions not annotated as promoters when determining if gene regulatory interactions were conserved after duplication. Improved gene models and epigenomic annotations in these regions will be crucial to fully understanding their function.

Gene annotation uncertainty also manifested when investigating proximal-to-distal transitions of regulatory elements. For example, many distal enhancers produce enhancer RNAs when actively regulating target genes (Sartorelli and Lauberth, 2020). Therefore, when investigating proximal-to-distal element transitions, it was not apparent whether the observed transcription at these elements was due to transcription from conserved promoter activity, or if these transcripts were enhancer RNAs. We also entertain a hypothesis that there may not always be a clear distinction between these types of transcription.

It is hard to disentangle between regulatory elements that were not effectively targeted with CRISPRi from orphaned regulatory elements that have no endogenous gene regulatory activity. It is therefore important not to conclude that an element does not have endogenous activity solely based on the lack of CRISPRi signal. Intrinsic differences in gRNA targeting efficiency exist *in vitro*, and we therefore aimed to test many guides per element. We also became more certain that a particular function was lost or reduced when we observed guides robustly affecting a different function of the same element. However, we are still limited by the current throughput of CRISPR-screening technology, both with respect to the number of guides that can be targeted to a particular element and the number of elements we can include in our screen. Additionally, single-cell CRISPRi screening experiments are not highly powered to detect regulatory element associations to lowly-expressed genes or regulatory elements which may have small, though phenotypically relevant (Richard et al., 2020; Muthuirulan et al., 2021), effect sizes.

## Supporting information

suppTable1

suppTable2

suppTable3

suppTable4

suppTable5

suppTable6

suppTable7

## Acknowledgments

We thank the Duke Human Vaccine Institute single-cell genomics core facility for their assistance. We thank Luke Bartelt, Raven Luo, Anushka Katikaneni, Hailey Napier and Natalie Dzikowski for critical feedback. We thank Yashodara Abeykoon for assistance with CRISPRi analysis. This research was supported by the Duke Whitehead Scholarship and the National Human Genome Research Institute (R35HG011332).

## Author Contributions

S.W. and C.B.L designed the study and wrote the paper. S.W. performed the experiments and analyses. C.B.L. supervised and funded research.

## Methods

### Self-alignment of hs1 human genome

To identify clusters of noncoding elements with high sequence identity in the human genome, we generated a self-alignment of the hs1 reference genome. First, the repeat-masked hs1 human reference genome was downloaded from the UCSC Genome Browser (Karolchik et al., 2012). We aligned every chromosome in the hs1 genome to itself and every other chromosome in the reference using LASTZ (Harris, 2007). The human.chimp.v2 scoring matrix, and alignment parameters for closely related species (O=600 E=150 T=2 M=254 K=4500 L=4500 Y=15000 C=0) were used for the alignment (Karolchik et al., 2012). In the case that a chromosome was aligned to itself, we used the *–self* parameter to filter out trivial one-to-one alignments. The generated alignments were output in LAV format and converted to AXT with *kentUtils:lavToAxt* (Kent et al., *2003). The resulting AXT files were concatenated and filtered to be larger than 200bp with gonomics:axTools* (Au et al., *2023)*.

#### Identifying putative regulatory elements

To identify putative regulatory elements in H9 hESCs and across *in vitro* cardiomyocyte differentiation, we downloaded unaligned H9 ATAC-seq sam files from *Liu et al*. (Table S6) with *SRA Toolkit:sam-dump* (https://github.com/ncbi/sra-tools/wiki/01-Downloading-SRA-Toolkit) (Liu et al., 2017). We converted the sam files to paired-end fastq with *samtools:sam2fq* and combined the fastq files across sample timepoints. Fastq files were then aligned to the hs1 genome using BWA MEM with default parameters (Li et al., 2009; Li and Durbin, 2009). Finally, we called open chromatin peaks with *macs3:callpeak* using default parameters for human genomes and generated a bed file corresponding to these peaks (Zhang et al., 2008).

### Clustering noncoding functional elements

We used the set of hs1 self-alignment AXTs and open chromatin peaks to identify clusters of noncoding elements in the hs1 human genome using *gonomics:liftWithAxt* and *gonomics:familiesFromLiftAndMerge*. Briefly, all open chromatin BED records were trimmed in length by 25% on both sides of the peak to limit regulatory elements to their core active bases using *gonomics:bedTrim*. The trimmed open chromatin peaks that were completely encompassed by an AXT alignment were lifted to their homologous location in the hs1 genome. Open chromatin peaks that had less that 70% identify to their lifted region were filtered out. Additionally, any lifted region that overlapped it’s original open chromatin peak (often the case in simple and tandem repeats) were filtered out. The whole set of elements, original open chromatin peaks and lifted open chromatin peaks, were merged together, keeping a record of which individual elements were joined. Any newly-joined element that did not contain an original open chromatin peak was given a unique name, “homologousElement_*N* “. All homologous elements were re-lifted through overlapping AXTs to draw connections between homologous elements.

We then created a graph-based representation of regulatory element families where elements (open chromatin or homologous elements) are represented by nodes and significant sequence identity between elements are represented by edges connecting nodes. In addition to the reported 1336 ESC clusters, our analysis framework identified 2648 extended clusters which contained only a single open chromatin region with many closed homologous elements. Since we are focused on how REFs can facilitate gene co-regulation, we excluded these 2648 single open element-clusters from our analysis. To visualize graphs of individual regulatory element families we used *gonomics:familiesFromLiftAndMerge* with the *- startNode* and *-igraph* options to generate a file compatible for visualization with a force-directed placement engine using the igraph R library (https://r.igraph.org/). Clustered elements were further visualized on a human T2T ideogram plot using the karyoploteR R library (Gel and Serra, 2017).

### Annotations of regulatory element families

We determined what gene annotations overlapped elements within ESC clusters. We generated 4 sets of elements for testing: all ESC open chromatin regions, open clustered elements, closed clustered elements, and the combined set of open and closed clustered elements. We next generated annotation files by filtering the hs1 NCBI Refseq GTF file, downloaded from the UCSC Genome Browser, for “transcripts,” “CDS,” “5UTR,” and, “3UTR.” We generated promoter annotations by taking 2kb upstream of the transcript start. We then used *kentUtils:overlapSelect* to sequentially overlap the annotations in the following order: promoters, 3’ UTR, 5’ UTR, CDS, transcripts. If an element overlapped an annotation it was classified as that annotation and removed from the next round of overlaps. If an element didn’t overlap exonic sequence (CDS), but overlapped a transcript, we annotated the element as “intronic.” If an element didn’t overlap any, we annotated that element as “intergenic.” We note that even though some clustered elements overlapped coding sequence, we expect these elements to also be strong candidates for having a noncoding function in ESCs. We performed this analysis for all four element sets.

To determine if relative proportions of proximal (promoters + 5’ UTR) vs distal (CDS + intron + 3’ UTR, intergenic) annotations differed between element sets, we performed hypergeometric tests. We first tested if proximal elements were enriched in open clustered elements vs the set of all ESC open chromatin peaks in R with the *phyper* function with the following settings: q = number of clustered proximal elements - 1; m = number of all ESC proximal elements; n = number of all ESC distal elements; k = number of open clustered elements, lower.tail = FALSE. We next tested if proximal elements were depleted in the closed subset vs the total set of open and closed clustered elements (all clustered elements). We again used the R function *phyper* with the following settings: q = number of closed proximal element;, m = number of proximal elements from all clustered elements; n = number of distal elements from all clustered elements; k = number of closed clustered elements; lower.tail = TRUE.

Next, to understand the mechanisms of how regulatory element families are created, we looked for enrichments of SDs in our regulatory element families dataset. We created two sets of noncoding clusters: the subset of open chromatin elements in clusters, and the subset of non-open chromatin elements in clusters. We then used *gonomics:overlapEnrichments* to calculate enrichments of hs1 annotated SDs (Vollger et al., 2022) in both clustered element sets.

### Gene expression analysis in ESCs and across in vitro CM differentiation

To integrate gene expression analysis we used time-point matched fragments per kilobase of transcript per million mapped reads (FPKM) RNA-seq data collected along with the ATAC-seq data from *Liu et al*. (Liu et al., *2017). We averaged FPKM across two replicates to generate a gene expression value for each gene. To compare REF8 regulatory element promoters to homologous inactive REF8 promoters, we used the R function t*.*test* with average FPKM values for the open and closed promoter gene sets. To determine which transcription factors were expressed at each timepoint across *in vitro* cardiomyocyte development, we subsetted the gene expression gene set by the transcription factor motifs we tested and created a 3 FPKM cutoff. Any genes with expression lower than this cutoff were considered not expressed at that timepoint.

### Identification of human-specific segmental duplications

Chimpanzee Oxford Nanopore (ONT) long read sequencing data was downloaded from the primate-T2T consortium GenomeArk browser (42basepairs.com/browse/s3/genomeark/species/Pan_troglodytes/mPanTro3/genomic_data/ont) (Yoo et al., 2025). Additionally we downloaded two human ONT sequencing datasets, one African individual and one Puerto Rican individual, from SRA (Table S6) (Shafin et al., 2020). For all three datasets, raw long-read ONT data was aligned to the hs1 human genome with *ngmlr* (Sedlazeck et al., *2018). Structural variation between the aligned human or chimpanzee ONT reads and the human reference were called with sniffles2* and output in VCF format (Smolka et al., 2024). We downloaded the dataset of SDs (Vollger et al., 2022) in hs1 coordinates from the UCSC Genome Browser. We first filtered the generated VCF files for deletions larger that 1kb in the aligned sequence (i.e. *insertions* in the reference relative to the aligned sequence). Human-specific SDs were called if the chimpanzee ONT-hs1 structural variants overlapped an hs1 SD by 50% and didn’t overlap a human polymorphic structural variant by more than 70% from the human ONT-hs1 alignments. Overlaps were generated using *kentUtils:overlapSelect* (Kent et al., *2003). Overlapping entries in the human-specific SD file were merged with gonomics:mergeBeds*.

### Enrichment of human-chimpanzee differentially expressed genes near human-specific SDs

To learn how human-specific SDs could be influencing gene expression, we analyzed species-specific differentially expressed genes from human-chimpanzee hybrid iPSCs from *Gokhman et al*., *2021* (Gokhman et al., 2021). *Differentially expressed genes were filtered for either cis-* or *trans-* regulatory differences. We then identified the number of species specific *cis-* and *trans-* regulated genes within 500kb of a human-specific SD. To determine if species specific *cis-* or *trans-* differentially expressed genes are enriched near human-specific SD, we randomly selected an number of genes equal to the number of true differentially expressed genes of that type (cis: 4280, trans: 3102) and determined how many were within 500kb of a human-specific SD. We iterated this process 1, 000, 000 times. We determined enrichment for each regulation type by comparing the number of true species-specific differentially expressed genes within 500kb of a human-specific SD to the matched random gene distribution. We generated a p-value by taking the number of trials where the randomly selected genes were greater than or equal to the true value, and dividing by the total number of trails.

### Human embryonic stem cell culture

H9 human embryonic stem cells (ESCs) (WiCell: WA09) obtained from WiCell (University of Wisconsin) were used for this study. H9s were grown in Matrigel-coated 6-well plates in mTeSR Plus media in a 37°C incubator with 5% CO_2_. Cells were passaged with ReleSR (STEMCELL) and grown in mTeSR Plus supplemented with 0.5x CloneR2 for 24 hours for standard passages.

### STARR-seq input library preparation

We tested 112 constructs from 11 different clusters in our STARR-seq assay. To generate each test construct, we identified 500bp from each annotated clustered element, and appended a unique 5 base-pair barcode to both sides of the test sequence for intra-family uniqueness. 20 negative control sequences were used: four previously published scrambled sequence controls (Mangan et al., 2022), 10 newly generated scrambled sequence controls (*gonomics:randSeq*), and 6 human genomic sequences with no chromatin accessibility in H9 ESCs (Table S7). STARR-seq constructs were ordered from Twist Biosciences with flanking cloning sites up and downstream of enhancer test sequence. Constructs were normalized and pooled before PCR amplification. The STARR-seq vector, *hSTARR-seq_ORI vector* was a gift from Alexander Stark (Addgene plasmid # 99296; http://n2t.net/addgene:99296; RRID:Addgene_99296) (Muerdter et al., 2018). First, *hSTARR-seq_ORI* was double digested with AgeI and SalI. Enhancer constructs were cloned into the STARR-seq vector with the NEB HiFi DNA Assembly kit. The cloned STARR-seq library was transformed into chemically competent cells (Gift from Ravi Karra) and after recovery, added to a 300 mL LB culture for overnight growth. The STARR-seq plasmid library was harvested with the Thermo Fisher GeneJet Endo-Free Plasmid Maxi Prep kit and the resulting library was concentrated with ethanol precipitation. The concentrated library was analyzed on a 1% agarose gel to assess overall plasmid quality. To assess construct representation in the resulting library, an input sequencing library was prepared with the NEB FS Ultra II DNA Library prep kit, with a 15-minute enzymatic fragmentation of the plasmid library. The input sequencing library was sequenced on the iSeq100 with 150 bp paired-end sequencing. We then quantified construct representation and generated an input normalization factor for each construct with *gonomics: starrSeqAnalysis inputSeq*.

### H9 hESC plasmid transfection

800,000 H9 ESCs per well were transfected with the Lonza 4D Nucleofector using the P3 Primary cell kit and the CB-150 transfection settings. In total, six transfections were completed, with 4ug of STARR-seq library and 1ug of pCAG-GFP per transfection. Transfected cells were plated in one well of a six-well plate coated with matrigel and cultured for 24 hours in mTesR plus supplemented with 1x CloneR2.

### STARR-seq output library creation

The output STARR-seq library preparation protocol was adapted from *Neumayr et al*. and *Johnson et al*. (Johnson et al., 2018; Neumayr et al., 2019). H9 ESCs were dissociated with Accutase (STEMCELL) and total RNA was extracted using the Qiagen RNeasy Plus Mini Prep kit with the gDNA eliminator column using two wells of a six-well plate per RNA prep, generating a total of 3 replicates for downstream analysis. 40 ug of total RNA per replicate was DNase Digested twice with the Thermo Fisher TURBO DNA-free kit. Total RNA was reverse transcribed for 2 hours at 50°C using Superscript IV VILO and a STARR-seq specific reverse transcription primer (SSRT). In tandem, a no RT control for each replicate was performed with nuclease-free water replacing the RT enzyme. Following reverse transcription, cDNA was cleaned up using 2x AMPure SPRI beads and eluted in 20uL nuclease-free water. The cDNA was then used for two rounds of STARR-seq library prep PCR. In PCR1, the primers “p7_seqPCR1” and “i5_PCR1” were used with the following cycling conditions for 20 cycles–denature: 98°C, annealing: 68°C, extension: 72°C. In PCR2, the primers “p7_seqPrimer” and “i5_PCR2_idX” were used to add a i5 sample index for sample multiplexing with the following cycling conditions for 15 cycles–denature: 98°C, annealing: 65°C, extension: 72°C. All primer sequences used can be found in Table S7. STARR-seq output libraries were pooled and sequenced at 100 pM on the iSeq100 with 20% PhiX using 150bp paired-end sequencing.

### STARR-seq analysis of regulatory element families

STARR-seq output libraries were de-multiplexed and converted to FASTQ with *bclConvert* (Version 4.1.5). A custom reference genome containing all test sequences was created using *gonomics: starrSeqAnalysis makeRef -byChrom -dualBx*. Genome index files were created with *STAR genomeGenerate* (version 2.7.11a). To protect against chimeric reporter transcripts being misaligned, Read 1 and Read 2 were aligned separately with *STAR*. The read 1 and read 2 flags were manually updated to reflect *first-in-pair* or *second-in-pair* status and read 1 and read 2 alignment files were concatenated and sorted by name with *samtools sort -n*. To quantify enhancer activity, all replicates were analyzed with *gonomics: starrSeqAnalysis bulkOutput -pe -dualBx –checkBx -bxHopping -zscore -inputNorm*. When a chimeric read was detected, a read in which reporter RNA from different REF elements within the same family annealed during PCR, we assigned a half-count to each element.

Negative control constructs with outlier normalized read counts were filtered out of the analysis by using Tukey’s method, or if they had zero raw counts in all three replicates. Fold enhancer activity was then calculated by taking a test construct’s normalized read-counts for a replicate and dividing it by the average of all negative control normalized read-counts from that replicate. Fold enhancer activities across replicates were averaged to generate a overall fold enhancer activity for each test construct. The Enhancer z-score statistic for each test construct is defined by taking the normalized read counts for that construct and comparing it to the mean and standard deviation of the negative controls in a replicate. Enhancer z-scores were combined across replicates for a given construct using Stauffer’s method to create a unified z-score. Test enhancer constructs were considered positive if they had a unified z-score higher than 2.32 (p < 0.01 in a one-tailed test).

### CRISPRi library design and cloning

To identify candidate enhancers in regulatory element families for CRISPRi screening, we downloaded raw FASTQs corresponding to ATAC-seq (Liu et al., 2017) and histone ChIP-seq data (H3k27ac, H3K3me1) (Battle et al., 2019) experiments in H9 ESCs from SRA (Table S6). FASTQs were aligned to the hs1 reference genome with BWA MEM and ChIP-seq alignment files for the same histone mark were merged with *samtools:merge*. Peaks were then called with *macs3:callPeak*. Putative enhancers were prioritized for selection by peak score for ATAC-seq and ChIP-seq signal, having human-specific members of regulatory element families, and visual inspection on the UCSC Genome Browser. A FASTA sequence corresponding to the epigenetic signal for each selected regulatory element was used for gRNA selection with the online CRISPick tool (Doench et al., 2016; Sanson et al., 2018). Briefly, we selected gRNAs, using hg38 for off target calculations for SpyoCas9, and got the top 10 ranked gRNAs per regulatory element. From those 10 gRNAs, we selected 6 gRNAs per element, prioritized by predicted activity and the gRNA’s ability to target additional members of the same cluster while not having genome-wide off target matches. Additionally, 24 non-targeting gRNAs were generated with *gonomics:randSeq* and cross referenced for having no homology to the human T2T genome. All guide sequences and targeting loci can be found in Table S7.

For all CRISPRi experiments the *pLV hU6-sgRNA hUbC-dCas9-KRAB-T2a-Puro* vector was used, a gift from Charles Gersbach (Addgene plasmid # 71236; http://n2t.net/addgene:71236; RRID:Addgene_71236) (Thakore et al., 2015).

CRISPR gRNA scaffolds were ordered from Twist Biosciences as single-stranded oligo pools with 2 gRNAs per oligo, flanked by BsmBI sites (Table S7). Oligos were made double stranded via PCR with primers to constant regions on either side of the oligos (Table S7). gRNAs were then cloned into the CRISPRi vector with Golden Gate cloning. Briefly, 0.12 pmol double-stranded gRNA scaffolds were added to 0.04 pmol uncut pLV hU6-sgRNA hUbC-dCas9-KRAB-T2a-Puro vector and incubated with Esp3I, T4 DNA Ligase, ATP, and rCutsmart for 2 hours for 37°C, 65°C for 15 minutes, 80°C for 10 minutes. Additional Esp3I was then added to digest any unassembled vector and the reaction was incubated at 37°C for 1 hour.

Concurrently, two 5 uL aliquots of the cloning reaction were transformed into 50uL of chemically competent Stbl3 cells (NEB) according to manufacturers protocols. After competent cell recovery, both transformation reactions were added to 300 mL LB with Carbenicillin and incubated overnight in a 37°C shaking incubator. The cells were harvested and the plasmid library was extracted with the Qiagen Midi Prep kit according to the manufacturer’s protocols.

To ensure even representation of CRISPRi gRNAs in the library, an Illumina sequencing library was prepped using the NEB Ultra II FS DNA Library Prep kit with enzymatic fragmentation of the plasmid library. The library was sequenced on the Illumina iSeq100 at 100 pmol with 20% PhiX with paired-end 150bp reads. Fastq reads were aligned to a custom reference containing all gRNA sequences with BWA MEM. PCR duplicates were removed from the alignment BAM file with *GATK:MarkDuplicates* (van der Auwera and O’Connor, 2020). gRNA abundance was quantified by aligned read-counts using *Gonomics:BedCountBam*.

### Lentivirus production

HEK293T cells (Duke Cell Culture Facility) were used for all lentivirus preparations. HEK293T cells were cultured in 15cm dishes in DMEM with high glucose, pyruvate, supplemented with 10% FBS and 1% Pen-Strep in a 37°C incubator with 5% CO_2_. During the lentivirus collection phase, DMEM was instead supplemented with 20% FBS and 1% Pen-Strep. HEK293T cells were transfected with a mixture of 7500 ng CRISPRi plasmid, 6570 ng PAX2 and 750 ng pMD2.G in OPTI-MEM with X-TremeGene. Virus-containing media was harvested at 24 and 48 hours. Lentivirus was concentrated with LentiX Concentrator and either used immediately or aliquoted and stored at −80°C.

### H9 hESC lentiviral transduction

H9 hESCs were transduced with lentivirus during a single-cell passaging using Accutase. 100, 000 Cells were transfected in mTeSR Plus media containing 10 ng/uL Polybrene and 1x CloneR2 (STEMCELL) and plated in a 6-well plate. After 24 hours, media was replaced with mTeSR Plus. After 48 hours, transduced cells were selected by supplementing mTeSR Plus media with 0.5 ng/uL Puromycin. After 10 days post-transduction, cells were harvested with Accutase and prepped for single-cell RNA-sequencing. Briefly, cells were washed with PBS and resuspended in PBS + 0.04% BSA.

### Single-Cell RNA-sequencing

Up to 10, 000 CRISPRi-transducted cells were captured in each of 2 lanes of the 10X Chromium Controller in collaboration with the Duke Human Vaccine Institute (DHVI). Gene expression and CRISPR feature libraries were prepared at the DHVI using the 5’ Gene Expression v3 Reagent kit with Feature Barcode Technology. The libraries were sequenced at the DHVI on the Illumina Nextseq2000 using a P3 flow cell.

### CRISPRi screening analysis

Raw BCL files were converted to FASTQ with *bclConvert* (Version 4.1.5) for each 10x lane separately. A custom hs1 10x reference was created with *cellranger:Mkref* (10x Genomics Cell Ranger v9.0.0) using the hs1 FASTA file and NCBI RefSeq GTF downloaded from the UCSC Genome Browser. *cellranger:Count* was run with the hs1 reference for each 10x lane separately. Each filtered count matrix output folder with feature barcode counts was analyzed in R using the package *Sceptre* (Barry et al., *2021). Briefly, gRNAs were assigned to cells with a 99% confidence threshold. We then tested for gRNA-gene expression associations within 200kb of targeted gRNA locations using the same hs1 GTF as in the cellranger count analysis. We used a one-sided test for repression of gene expression and either the union* or *singleton* gRNA integration strategy to test for association. Both of these analysis frameworks are useful. In our union analysis, we tested all element-gene pairs within 200kb that passed the default quality control thresholds. Element-gene association tests were further filtered out if either the treatment set of cells (all cells that received a targeting gRNA to that element) or control set of cells (all cells that did not receive a targeting gRNA to that element) had less than seven cells that had non-zero gene expression counts for the tested gene. For clarity in the results, we defined the expression-filtered genes as “not expressed.” Due to intrinsic differential activities of gRNAs targeting the same element, significant individual gRNA-gene expression associations can be washed out in a union analysis if the other gRNAs targeting the same element have low repressive activity. Therefore, the singleton analysis can identify gRNA-expression pairs that may be lost in the union analysis. In our singleton analysis, since statistical power is generally lower, we raised our gene expression to only test genes with higher expression. We raised the treatment and control non-zero cells filters to 20 and 3000, respectively.

In the case that a gRNA targeted more than location in the genome, gRNA-gene expression associations were tested for all locations. We considered only perfect gRNA sequence matches to the hs1 genome sequence as a proper targeting gRNA loci. It is possible that there were “off target” effects of gRNAs within REF enhancer paralogs with higher divergence, but we did not test for these associations in our analysis. Additionally, we input the top 5 principle components from a Seurat (v5.2.1) analysis as covariates to the analysis formula (Hao et al., 2021). Per sceptre’s default settings, raw p-values were corrected with the Benjamini-Hochberg method with a false discovery rate (FDR) of 0.1. We additionally filtered for significant gRNA-gene pairs to have a log2-fold change beyond −0.1.

### Inferring alternative REF1 transcripts

Based on the alignment of H9 ESC RNA-seq from the CRISPRi experiment to the human T2T reference, we hypothesized that some genes with REF1 element promoters were either mis-annotated or the primary isoforms were missing from the reference. We used BLAT (Kent et al., 2003) to map exons from *LOC124906734*, the REF1-proximal transcript that best matched the RNA-seq alignments, to other locations in the human T2T reference genome. We used the BLAT-mapped exons to infer transcripts that may better reflect the mRNA that exists in H9 ESCs. We annotated these inferred transcripts in figures with a lighter green color than the standard NCBI transcript set.

### Addition of hg38-lifted transcripts to human T2T gene set

We observed that some putative transcripts of interest within SDs were not in the human T2T gene set. Therefore, we added putative transcripts to human T2T gene set and re-ran the CRISPRi analysis. Briefly, putative transcripts were either selected from the “CAT/Liftoff Gene” track on the human T2T UCSC genome browser (Fiddes et al., 2018a) or lifted with BLAT (Kent et al., 2003). *LINC03072* was mapped with BLAT from the hg38 human genome reference. Transcripts from the CAT/Liftoff gene set were renamed as the following:

| CAT/Liftoff name | Name in text |
| --- | --- |
| AC090114.3-201 | TLK2-like-chr7 |
| AC004923.1-201 | LOC422-chr11 |

With the genes added to the human T2T gene set, we re-ran the cellranger pipeline and a union sceptre analysis as previously described.

### Visualization of UCSC Genome Browser data

To provide context to regulatory elements for which we had CRISPRi and STARR-seq data, we generated visuals adapted from the UCSC genome browser (Karolchik et al., 2012). Briefly, we loaded the tracks we wanted to display and saved the browser viewing window as an image. In Adobe Illustrator we used the image trace function to generate vectors of genomic data tracks (ie: ATAC-seq or RNA-seq), and manually created gene models for genes in the NCBI database. Coordinates are from the human T2T (hs1) assembly unless otherwise noted.

### Alignment of paralogous SD

We aimed to understand the sequence conservation and alignment of SDs on chromosome 4 and chromosome 11, which harbored elements from REF1 and REF3. We defined the SD breakpoints using the UCSC Genome Browser and the SD track (Numanagić et al., 2018; Vollger et al., 2022). We downloaded the resulting fasta sequences, and since the two paralogous SD were inverted compared to one another, we took the reverse compliment of the chromosome 4 SD with *gonomics:faFormat -revComp* (Au et al., 2023). We aligned the resulting sequences with lastz (Harris, 2007). Specifically we used the default scoring matrix and the same command line options as for the whole-genome self alignment and output the alignment in PAF format. We filtered the resulting PAF alignments to be larger than 3000bp with the R library *SVbyEye* and visualized the alignment with the *plotMiro* function (Porubsky et al., 2025).

### Evolutionary relationships of REF4/5-containing SDs

To identify the primate-syntenic REF4/5 elements on chromosome 1, we used BLAT (Kent et al., 2003) to map a representative REF4/5 sequence to each primate genome (chimpanzee, gorilla, orangutan) (Yoo et al., 2025). To further determine copy number of REF4/5 elements in the respective primate genomes, we used BLAT to map the respective primate homolog of REF4/5 to the complete primate genome. We used a 70% sequence identity filter to call significant alignments. To determine the source copies of the human-specific REF4/5-containing SDs, we analyzed the human SD track on the UCSC genome browser (Karolchik et al., 2012; Numanagić et al., 2018; Vollger et al., 2022) to identify human loci with the longest and highest-scoring contiguous alignments between SD copies in the human genome.

### Transcription factor binding motif enrichment

Position Frequency Matrices (PFMs) for core vertebrate transcription factors were downloaded from the JASPAR database (https://jaspar2020.genereg.net/downloads/) Fasta sequences for time point-specific families were obtained by running *gonomics: bedToFasta* and converting soft-masked bases in the fasta file to uppercase with *gonomics: faFormat -toUpper*. Transcription factor binding motif identification was then performed with *gonomics:tfMatch* with default settings. To perform enrichment analysis, we used the R function *pbinom* with *lower*.*tail = FALSE* to model a binomial distribution for the probability of obtaining greater than or equal to the number of observed motif matches for a given transcription factor in the test set compared to the control set. We generated an initial probability of observing a motif match in the test set by taking the length (in base pairs) of all sequences in the test set and dividing by the total length of all TP-family sequences. We restricted enrichment calculations to transcription factors that are expressed at relevant timepoints, and transcription factors that had minimum motif matches among all sequences to reach Bonferroni-adjusted significance (p < 0.05) if all matches were to be in the test set. The number of motif matches required for testing in H9, cardiac mesoderm, and cardiomyocytes were 6, 19, and 4, respectively. After p-values were generated, we performed Bonferroni correction to generate adjusted p-values. We additionally calculated fold-enrichment by first modeling the expected number of motif matches in the test. This is done by taking the baseline probability of observing a motif match in the test set and multiplying by the total number of observed motif matches among all sequences from the test and control sets. Fold-enrichment can then be obtained by dividing the observed number of motif matches in the test set by the expected number of motif matches in the test set. To visualize the significant transcription factor motif enrichments, we plotted fold enrichment with the *pheatmap* R library https://github.com/raivokolde/pheatmap.

### Enrichment of ENCODE cell types

First, all human ENCODE ATAC-seq and DNase-seq fastq files with read length longer than 75 nucleotides were downloaded. When multiple fastq files came from the same biosample classification, or a single sample had multiple replicates, we kept ATAC-seq over DNase-seq, kept longer read lengths over shorter, and kept samples with deeper read depth until we had a single fastq dataset for each biosample (Table S6). The remaining 154 fastq files were aligned to the human T2T genome and called open chromatin peaks as previously described.

To determine what cell types had a high degree of noncoding clusters, and had a high degree of human-specific cluster expansion, we used *gonomics:overlapEnrichments* with each ENCODE peak set and human T2T loci with self-paralogy or human-specific SDs, respectively. The resulting p-values were Bonferroni-corrected. We plotted fold enrichment with −log10(adjusted p-value) and in the case of multiple closely-related biosamples (e.g. multiple B-cell types) only the sample with the highest fold enrichment was plotted for clarity.

## Data and Code Availability

Supplementary material is available at *Molecular Biology and Evolution* online.

Software written for this manuscript were implemented as a part of Gonomics, an ongoing effort to develop an open-source genomics platform in the Go programming language (golang). Gonomics can be accessed at https://github.com/vertgenlab/gonomics.

Additional software, as well as raw and analyzed datasets, including browser tracks, sequencing files, alignments, and figure generation pipelines are available on our lab website at https://www.vertgenlab.org and in GEO under bioproject PRJNA1367149.

## Supplementary Figures

Supplementary figures to accompany the main text.

**Figure S1.**
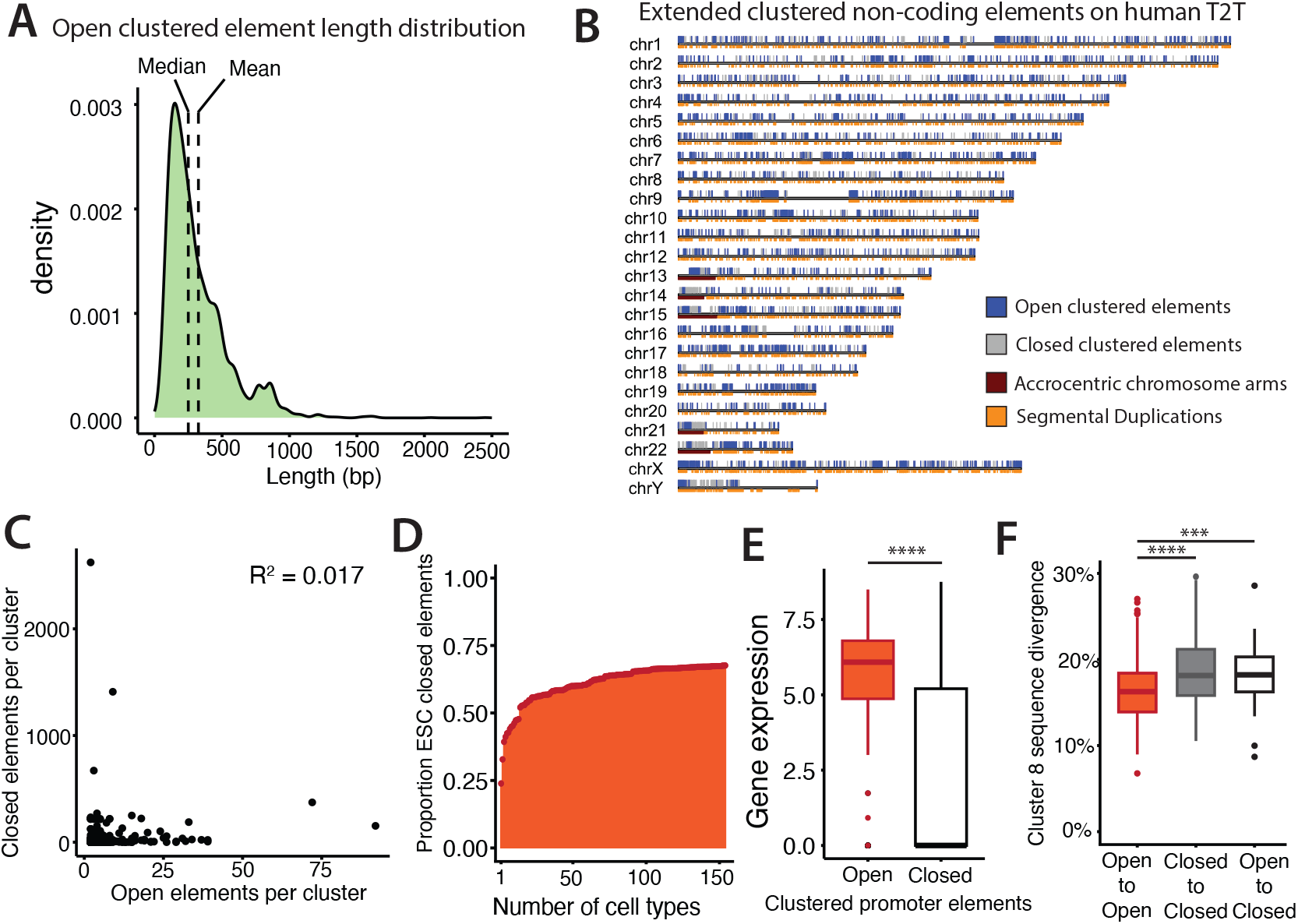
Open and closed clustered elements harbor differences in ESCs. **(A)** Location of embryonic stem cell (ESC) open clustered elements, closed clustered elements, segmental duplications and acrocentric chromosome arms in the human T2T reference genome. **(B)** Size distributions of open ESC clustered elements Eleven elements larger than 2500 bp were filtered out for ease of visualization. **(C)** Correlation of number of open and closed ESC elements per cluster. **(D)** Proportion of ESC closed elements that have accessible chromatin in other ENCODE cell types as increasing cell types are considered. **(E)** Comparison of gene expression between gene with open vs closed ESC promoters. Y-axis values reflect Log10-transformed fragments per kilobase plus a pseudocount. (Wilcoxon; ****: p *<* 0.0001). **(F)** Percent divergence between extended Cluster 8 elements, comparing sequence divergence between open elements, between open and closed elements, and between closed elements. (t-test; ***: p *<* 0.001, ****: p *<* 0.0001)

**Figure S2.**
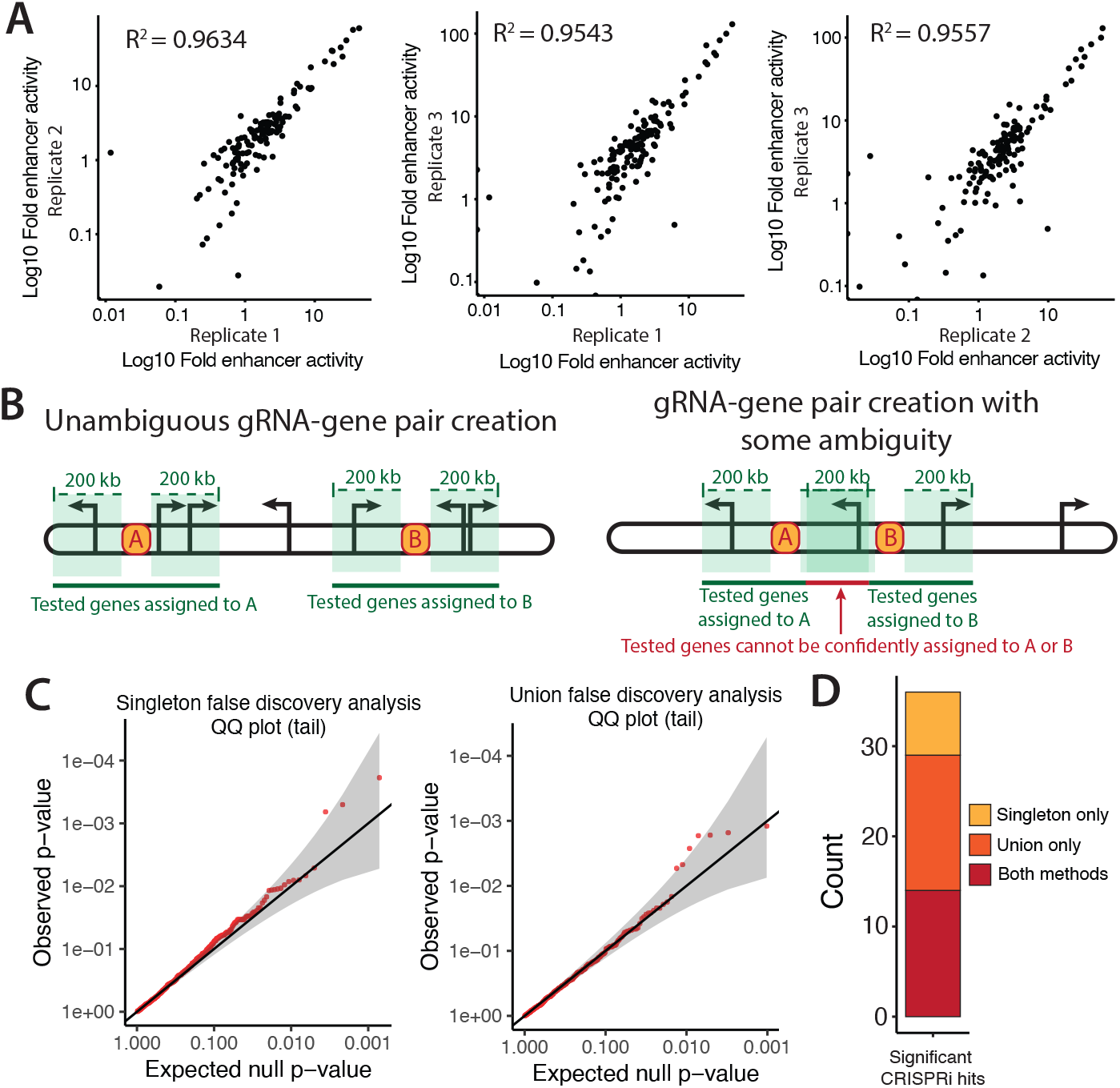
Functional data analysis quality. **(A)** Correlation of three STARR-seq replicates performed in H9 ESCs. Axes are Log10 transformed **(B)** Representation of gRNA-gene pair creation for association testing by sceptre with a 200kb distance filter, where elements “A” and “B” are paralogous in a cluster. Arrows denote location of genes in the model. **(C)** Quality-control false discovery QQ plots produced by sceptre for both singleton and union analyses. No false positives were detected in either analysis. **(D)** Number of significant clustered element CRISPRi hits that were found in the singleton analysis, the union analysis or both analysis methods.

**Figure S3.**
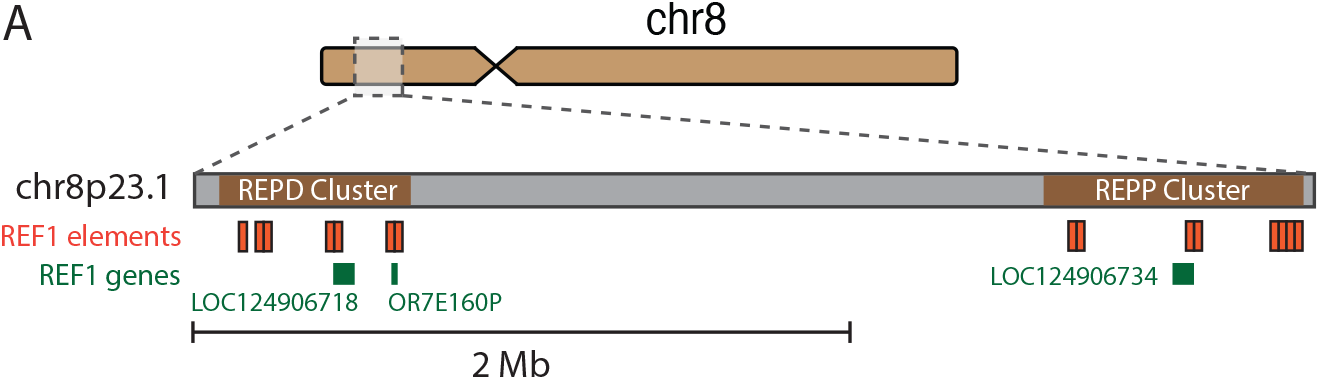
Chromosome 8 SD clusters harbor REF1 promoters. (**A)** Graphical representation of REF1 elements within segmental duplication clusters on chromosome 8p23.1.

**Figure S4.**
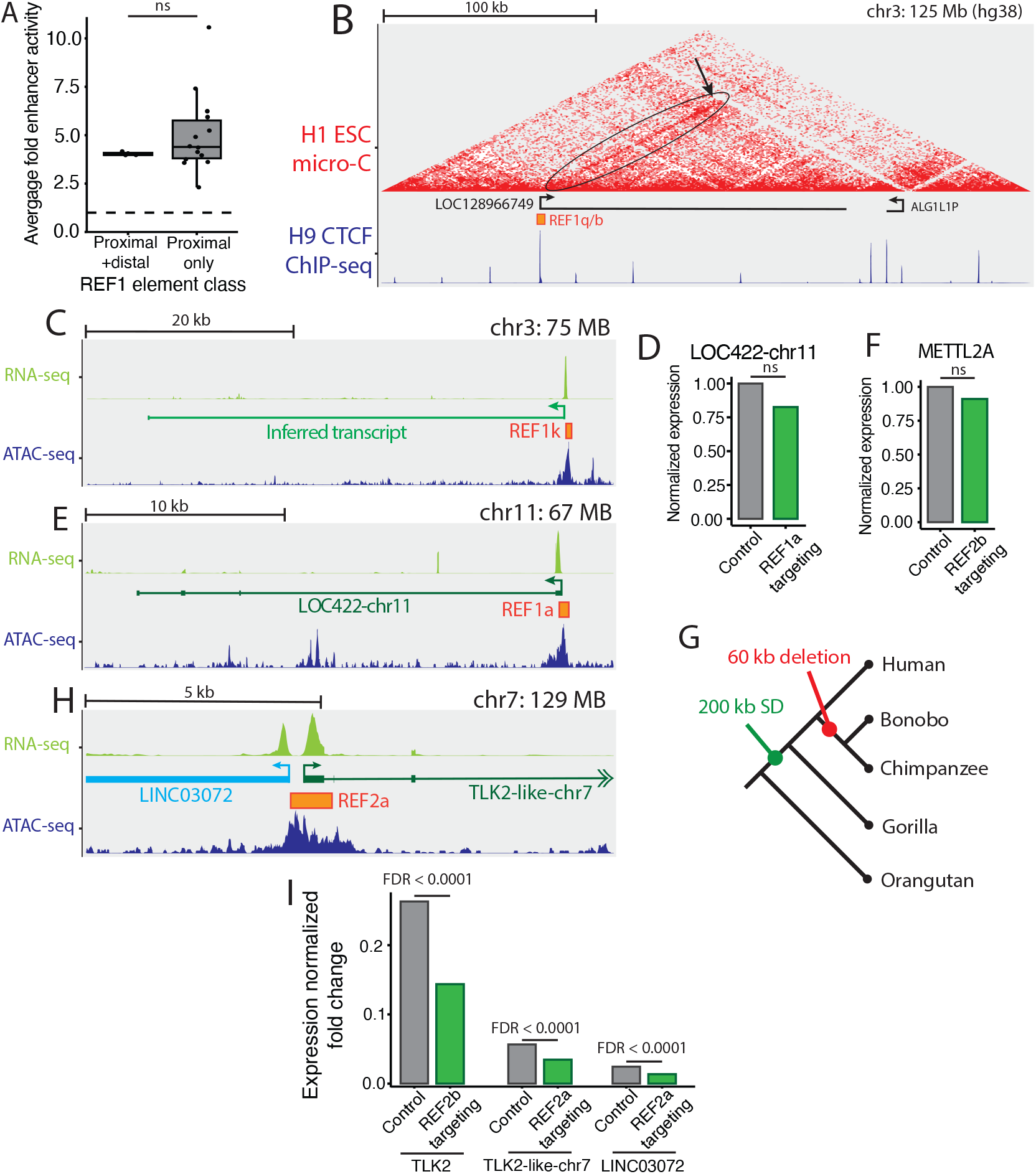
Element class flexibility associated data. **(A)** Comparison of enhancer activity between CRISPRi-targeted REF1 elements that had proximal and distal regulatory activity compared to REF1 elements that had proximal activity only, measured by STARR-seq (Wilcoxon test, ns: p *>* 0.05). **(B)** 3D genome organization of the *LOC128966749*-*ALG1L1P* promoter-promoter interaction visualized on the hg38 genome assembly. The oval shows the stripe domain facilitated by the *LOC128966749* promoter, and the arrow shows the interaction with the *ALG1L1P* promoter. **(C)** Genomic context for REF1k. A transcription start site was inferred based on 5’-capture RNA-seq from ESCs and a gene model lifted from the hg38 assembly. **(D)** Normalized *LOC422-chr11* expression upon repression of REF1a with CRISPRi (ns: Benjamini-Hochberg FDR *>* 0.1) **(E)** Genomic context for REF1a. *LOC422-chr11* was lifted from the hg38 reference. **(F)** Normalized *METTL2A* expression upon repression of REF2b with CRISPRi (ns: Benjamini-Hochberg FDR *>* 0.1). **(G)** Primate phylogeny with arbitrary branch lengths showing the duplication history of the segmental duplication that created REF2a. **(H)** Genomic context of REF2a. Genes *TLK-chr7* and *LINC03072* lifted onto the human T2T assembly from hg38. **(I)** Fold-change gene expression plots showing conserved regulatory interactions when REF2 elements are repressed with CRISPRi. Control and targeting gene expression bars are scaled by gene expression in all cells that did not receive a targeting gRNA, as calculated by Seurat (Benjamini-Hochberg FDR *<* 0.1).

**Figure S5.**
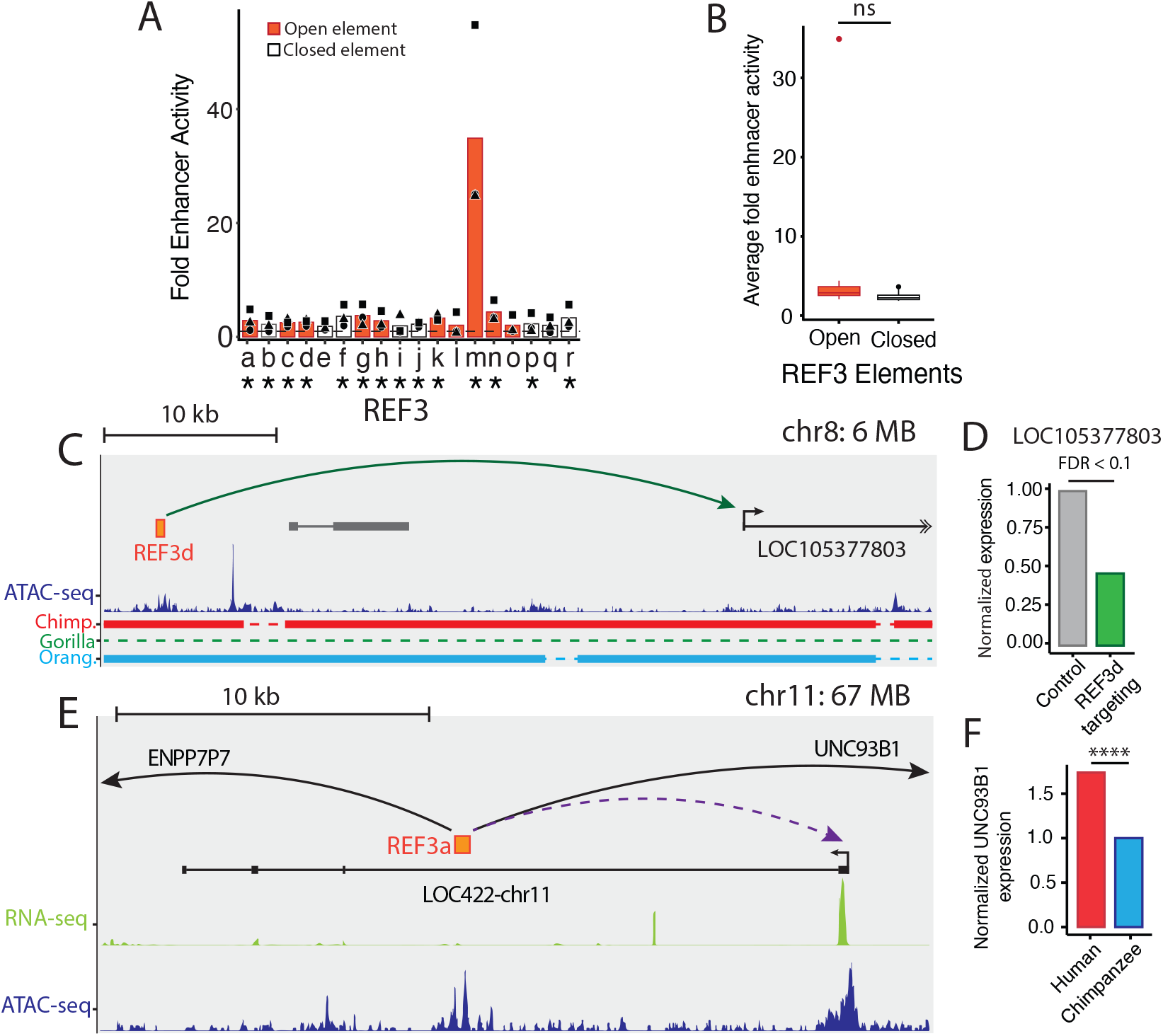
REF3 rewiring associated data. **(A)** Fold enhancer activity of REF3 elements over negative controls. Data point shape denotes replicate number. Dotted line is the average of negative controls (Stauffer’s method, *: p *<* 0.01). **(B)** Comparison of fold enhancer activity in REF3 between open and closed elements (Wilcoxon; ns: p *>* 0.5). **(C)** Genomic context for REF3d. Curved arrows represent a distal regulatory connection. **(D)** Normalized *LOC105377803* expression upon CRISPRi repression of REF3a (Benjamini-Hochberg FDR *>* 0.1) **(E)** Genomic context of REF3a. The *LOC422-chr11* gene model was lifted onto human T2T from hg38. Curved arrows represent distal regulatory interactions. **(F)** Human *UNC93B1* expression normalized to chimpanzee *UNC93B1* expression. Data from *Gokhman et. al*., *2021*, using human-chimpanzee allotetraploid cells (Differential expression test; p *<* 0.0001: ****).

**Figure S6.**
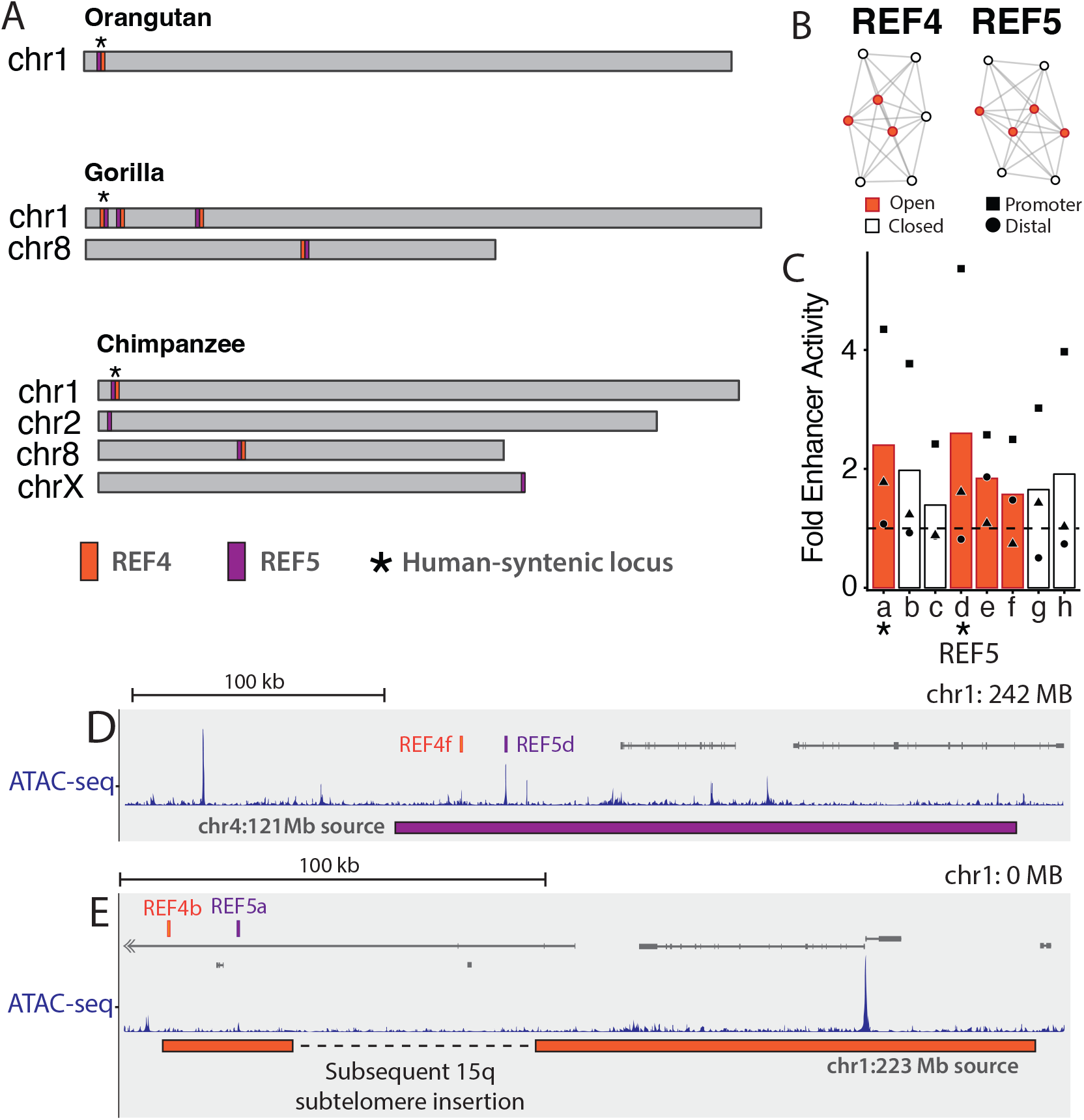
Lineage-specific duplications of REF4 and REF5. **(A)** Locations of REF4 and REF5 elements in T2T primate genomes. **(B)** REF4 and REF5 visualized with edge lengths corresponding to percent divergence between noncoding element nodes. **(C)** Fold enhancer activity of REF5 elements over negative controls, measured by STARR-seq. Data point shape denotes replicate number. Dotted line is the average of negative controls (Stauffer’s method, *: p *<* 0.01). **(D-E)** Genomic context showing the source sequence for human-specific SDs containing REF4/5 elements.

## References

Ali, A., Han, K., and Liang, P., 2021. Role of transposable elements in gene regulation in the human genome. Life, 11(2):118.

Allocco, D. J., Kohane, I. S., and Butte, A. J., 2004. Quantifying the relationship between co-expression, coregulation and gene function. BMC bioinformatics, 5:1–10.

Andersson, R. and Sandelin, A., 2020. Determinants of enhancer and promoter activities of regulatory elements. Nature Reviews Genetics, 21(2):71–87.

Anzai, T., Shiina, T., Kimura, N., Yanagiya, K., Kohara, S., Shigenari, A., Yamagata, T., Kulski, J. K., Naruse, T. K., Fujimori, Y., et al., 2003. Comparative sequencing of human and chimpanzee mhc class i regions unveils insertions/deletions as the major path to genomic divergence. Proceedings of the National Academy of Sciences, 100(13):7708–7713.

Arnold, C. D., Gerlach, D., Stelzer, C., Boryń, Ł. M., Rath, M., and Stark, A., 2013. Genome-wide quantitative enhancer activity maps identified by starr-seq. Science, 339(6123):1074–1077.

Au, E. H., Fauci, C., Luo, Y., Mangan, R. J., Snellings, D. A., Shoben, C. R., Weaver, S., Simpson, S. K., and Lowe, C. B., 2023. Gonomics: uniting high performance and readability for genomics with go. Bioinformatics, 39(8):btad516.

Barakat, T. S., Halbritter, F., Zhang, M., Rendeiro, A. F., Perenthaler, E., Bock, C., and Chambers, I., 2018. Functional dissection of the enhancer repertoire in human embryonic stem cells. Cell stem cell, 23(2):276–288.

Barber, J. C., Maloney, V. K., Huang, S., Bunyan, D. J., Cresswell, L., Kinning, E., Benson, A., Cheetham, T., Wyllie, J., Lynch, S. A., et al., 2008. 8p23. 1 duplication syndrome; a novel genomic condition with unexpected complexity revealed by array cgh. European journal of human genetics, 16(1):18–27.

Barry, T., Roeder, K., and Katsevich, E., 2024. Exponential family measurement error models for single-cell crispr screens. Biostatistics, 25(4):1254–1272.

Barry, T., Wang, X., Morris, J. A., Roeder, K., and Katsevich, E., 2021. Sceptre improves calibration and sensitivity in single-cell crispr screen analysis. Genome biology, 22(1):344.

Battle, S. L., Jayavelu, N. D., Azad, R. N., Hesson, J., Ahmed, F. N., Overbey, E. G., Zoller, J. A., Mathieu, J., Ruohola-Baker, H., Ware, C. B., et al., 2019. Enhancer chromatin and 3d genome architecture changes from naive to primed human embryonic stem cell states. Stem cell reports, 12(5):1129– 1144.

Baur, J. A., Zou, Y., Shay, J. W., and Wright, W. E., 2001. Telomere position effect in human cells. Science, 292(5524):2075–2077.

Beisaw, A., Kuenne, C., Guenther, S., Dallmann, J., Wu, C.-C., Bentsen, M., Looso, M., and Stainier, D. Y., 2020. Ap-1 contributes to chromatin accessibility to promote sarcomere disassembly and cardiomyocyte protrusion during zebrafish heart regeneration. Circulation research, 126(12):1760– 1778.

Bejerano, G., Haussler, D., and Blanchette, M., 2004. Into the heart of darkness: large-scale clustering of human non-coding dna. Bioinformatics, 20(suppl_1):i40–i48.

Bejerano, G., Lowe, C. B., Ahituv, N., King, B., Siepel, A., Salama, S. R., Rubin, E. M., James Kent, W., and Haussler, D., 2006. A distal enhancer and an ultraconserved exon are derived from a novel retroposon. Nature, 441(7089):87–90.

Bejjani, F., Evanno, E., Zibara, K., Piechaczyk, M., and Jariel-Encontre, I., 2019. The ap-1 transcriptional complex: Local switch or remote command? Biochimica et Biophysica Acta (BBA)-Reviews on Cancer, 1872(1):11–23.

Benton, M. L., Abraham, A., LaBella, A. L., Abbot, P., Rokas, A., and Capra, J. A., 2021. The influence of evolutionary history on human health and disease. Nature Reviews Genetics, 22(5):269–283.

Blekhman, R., Oshlack, A., and Gilad, Y., 2009. Segmental duplications contribute to gene expression differences between humans and chimpanzees. Genetics, 182(2):627– 630.

Bosch, N., Cáceres, M., Cardone, M. F., Carreras, A., Ballana, E., Rocchi, M., Armengol, L., and Estivill, X., 2007. Characterization and evolution of the novel gene family fam90a in primates originated by multiple duplication and rearrangement events. Human molecular genetics, 16(21):2572–2582.

Britten, R. J. and Davidson, E. H., 1971. Repetitive and nonrepetitive dna sequences and a speculation on the origins of evolutionary novelty. The Quarterly review of biology, 46(2):111–138.

Britten, R. J. and Kohne, D. E., 1968. Repeated sequences in dna: hundreds of thousands of copies of dna sequences have been incorporated into the genomes of higher organisms. Science, 161(3841):529–540.

Caglayan, E., Ayhan, F., Liu, Y., Vollmer, R. M., Oh, E., Sherwood, C. C., Preuss, T. M., Yi, S. V., and Konopka, G., 2023. Molecular features driving cellular complexity of human brain evolution. Nature, 620(7972):145–153.

Chuong, E. B., Elde, N. C., and Feschotte, C., 2016. Regulatory evolution of innate immunity through co-option of endogenous retroviruses. Science, 351(6277):1083–1087.

Chuong, E. B., Elde, N. C., and Feschotte, C., 2017. Regulatory activities of transposable elements: from conflicts to benefits. Nature Reviews Genetics, 18(2):71–86.

Cooper, G. M., Coe, B. P., Girirajan, S., Rosenfeld, J. A., Vu, T. H., Baker, C., Williams, C., Stalker, H., Hamid, R., Hannig, V., et al., 2011. A copy number variation morbidity map of developmental delay. Nature genetics, 43(9):838–846.

Dennis, M. Y. and Eichler, E. E., 2016. Human adaptation and evolution by segmental duplication. Current opinion in genetics & development, 41:44–52.

Dennis, M. Y., Nuttle, X., Sudmant, P. H., Antonacci, F., Graves, T. A., Nefedov, M., Rosenfeld, J. A., Sajjadian, S., Malig, M., Kotkiewicz, H., et al., 2012. Evolution of human-specific neural srgap2 genes by incomplete segmental duplication. Cell, 149(4):912–922.

Doench, J. G., Fusi, N., Sullender, M., Hegde, M., Vaimberg, E. W., Donovan, K. F., Smith, I., Tothova, Z., Wilen, C., Orchard, R., et al., 2016. Optimized sgrna design to maximize activity and minimize off-target effects of crisprcas9. Nature biotechnology, 34(2):184–191.

Dougherty, M. L., Underwood, J. G., Nelson, B. J., Tseng, E., Munson, K. M., Penn, O., Nowakowski, T. J., Pollen, A. A., and Eichler, E. E., 2018. Transcriptional fates of human-specific segmental duplications in brain. Genome research, 28(10):1566–1576.

Duan, Y.-Y., Chen, X.-F., Zhu, R.-J., Jia, Y.-Y., Huang, X.-T., Zhang, M., Yang, N., Dong, S.-S., Zeng, M., Feng, Z., et al., 2023. High-throughput functional dissection of noncoding snps with biased allelic enhancer activity for insulin resistance-relevant phenotypes. The American Journal of Human Genetics, 110(8):1266–1288.

ENCODE Project Consortium, 2012. An integrated encyclopedia of dna elements in the human genome. Nature, 489(7414):57.

Falker-Gieske, C., 2023. Transcriptome driven discovery of novel candidate genes for human neurological disorders in the telomer-to-telomer genome assembly era. Human Genomics, 17(1):94.

Feschotte, C., 2008. Transposable elements and the evolution of regulatory networks. Nature Reviews Genetics, 9(5):397– 405.

Fiddes, I. T., Armstrong, J., Diekhans, M., Nachtweide, S., Kronenberg, Z. N., Underwood, J. G., Gordon, D., Earl, D., Keane, T., Eichler, E. E., et al., 2018a. Comparative annotation toolkit (cat)—simultaneous clade and personal genome annotation. Genome research, 28(7):1029–1038.

Fiddes, I. T., Lodewijk, G. A., Mooring, M., Bosworth, C. M., Ewing, A. D., Mantalas, G. L., Novak, A. M., van den Bout, A., Bishara, A., Rosenkrantz, J. L., et al., 2018b. Human-specific notch2nl genes affect notch signaling and cortical neurogenesis. Cell, 173(6):1356–1369.

Finlayson-Short, L., Davey, C. G., and Harrison, B. J., 2020. Neural correlates of integrated self and social processing. Social cognitive and affective neuroscience, 15(9):941–949.

Foster, B. L., Koslov, S. R., Aponik-Gremillion, L., Monko, M. E., Hayden, B. Y., and Heilbronner, S. R., 2023. A tripartite view of the posterior cingulate cortex. Nature Reviews Neuroscience, 24(3):173–189.

Fraimovitch, E. and Hagai, T., 2023. Promoter evolution of mammalian gene duplicates. BMC biology, 21(1):80.

Franke, M., Ibrahim, D. M., Andrey, G., Schwarzer, W., Heinrich, V., Schöpflin, R., Kraft, K., Kempfer, R., Jerković, I., Chan, W.-L., et al., 2016. Formation of new chromatin domains determines pathogenicity of genomic duplications. Nature, 538(7624):265–269.

Gasperini, M., Hill, A. J., McFaline-Figueroa, J. L., Martin, B., Kim, S., Zhang, M. D., Jackson, D., Leith, A., Schreiber, J., Noble, W. S., et al., 2019. A genome-wide framework for mapping gene regulation via cellular genetic screens. Cell, 176(1):377–390.

Gel, B. and Serra, E., 2017. karyoploter: an r/bioconductor package to plot customizable genomes displaying arbitrary data. Bioinformatics, 33(19):3088–3090.

Ghafouri-Fard, S., Harsij, A., Hussen, B. M., Taheri, M., and Ayatollahi, S. A., 2023. A review on the role of snhg8 in human disorders. Pathology-Research and Practice, 245:154458.

Gokhman, D., Agoglia, R. M., Kinnebrew, M., Gordon, W., Sun, D., Bajpai, V. K., Naqvi, S., Chen, C., Chan, A., Chen, C., et al., 2021. Human–chimpanzee fused cells reveal cisregulatory divergence underlying skeletal evolution. Nature genetics, 53(4):467–476.

Grandi, N. and Tramontano, E., 2018. Human endogenous retroviruses are ancient acquired elements still shaping innate immune responses. Frontiers in immunology, 9:2039.

Guterstam, A., Björnsdotter, M., Gentile, G., and Ehrsson, H. H., 2015. Posterior cingulate cortex integrates the senses of self-location and body ownership. Current Biology, 25(11):1416–1425.

Hao, Y., Hao, S., Andersen-Nissen, E., Mauck, W. M., Zheng, S., Butler, A., Lee, M. J., Wilk, A. J., Darby, C., Zager, M., et al., 2021. Integrated analysis of multimodal single-cell data. Cell, 184(13):3573–3587.

Hardison, R. C., 2012. Evolution of hemoglobin and its genes. Cold Spring Harbor perspectives in medicine, 2(12):a011627.

Harris, R. S., 2007. Improved pairwise alignment of genomic DNA. The Pennsylvania State University.

He, P., Zhang, C., Ji, Y., Ge, M.-K., Yu, Y., Zhang, N., Yang, S., Yu, J.-X., Shen, S.-M., and Chen, G.-Q., et al., 2022. Epithelial cells-enriched lncrna snhg8 regulates chromatin condensation by binding to histone h1s. Cell Death & Differentiation, 29(8):1569–1581.

Hong, J.-W., Hendrix, D. A., and Levine, M. S., 2008. Shadow enhancers as a source of evolutionary novelty. Science, 321(5894):1314–1314.

Horton, I., Kelly, C. J., Dziulko, A., Simpson, D. M., and Chuong, E. B., 2023. Mouse b2 sine elements function as ifn-inducible enhancers. elife, 12:e82617.

Huo, Y., Cao, K., Kou, B., Chai, M., Dou, S., Chen, D., Shi, Y., and Liu, X., 2023. Tp53bp2: Roles in suppressing tumorigenesis and therapeutic opportunities. Genes & Diseases, 10(5):1982–1993.

Jeong, H., Dishuck, P. C., Yoo, D., Harvey, W. T., Munson, K. M., Lewis, A. P., Kordosky, J., Garcia, G. H., (HGSVC), H. G. S. V. C., Yilmaz, F., et al., 2025. Structural polymorphism and diversity of human segmental duplications. Nature genetics, :1–12.

Johnson, G. D., Barrera, A., McDowell, I. C., D’Ippolito, A. M., Majoros, W. H., Vockley, C. M., Wang, X., Allen, A. S., and Reddy, T. E., 2018. Human genome-wide measurement of drug-responsive regulatory activity. Nature communications, 9(1):5317.

Ju, X.-C., Hou, Q.-Q., Sheng, A.-L., Wu, K.-Y., Zhou, Y., Jin, Y., Wen, T., Yang, Z., Wang, X., and Luo, Z.-G., et al., 2016. The hominoid-specific gene tbc1d3 promotes generation of basal neural progenitors and induces cortical folding in mice. Elife, 5:e18197.

Karolchik, D., Hinrichs, A. S., and Kent, W. J., 2012. The ucsc genome browser. Current Protocols in Bioinformatics, 40(1).

Kent, W. J., Baertsch, R., Hinrichs, A., Miller, W., and Haussler, D., 2003. Evolution’s cauldron: duplication, deletion, and rearrangement in the mouse and human genomes. Proceedings of the National Academy of Sciences, 100(20):11484–11489.

Keough, K. C., Whalen, S., Inoue, F., Przytycki, P. F., Fair, T., Deng, C., Steyert, M., Ryu, H., Lindblad-Toh, K., Karlsson, E., et al., 2023. Three-dimensional genome rewiring in loci with human accelerated regions. Science, 380(6643):eabm1696.

Kim, G.-J., Sock, E., Buchberger, A., Just, W., Denzer, F., Hoepffner, W., German, J., Cole, T., Mann, J., Seguin, J. H., et al., 2015. Copy number variation of two separate regulatory regions upstream of sox9 causes isolated 46, xy or 46, xx disorder of sex development. Journal of Medical Genetics, 52(4):240–247.

Krietenstein, N., Abraham, S., Venev, S. V., Abdennur, N., Gibcus, J., Hsieh, T.-H. S., Parsi, K. M., Yang, L., Maehr, R., Mirny, L. A., et al., 2020. Ultrastructural details of mammalian chromosome architecture. Molecular cell, 78(3):554–565.

Leech, R. and Sharp, D. J., 2014. The role of the posterior cingulate cortex in cognition and disease. Brain, 137(1):12– 32.

Li, H. and Durbin, R., 2009. Fast and accurate short read alignment with burrows–wheeler transform. bioinformatics, 25(14):1754–1760.

Li, H., Handsaker, B., Wysoker, A., Fennell, T., Ruan, J., Homer, N., Marth, G., Abecasis, G., Durbin, R., and Subgroup, . G. P. D. P., et al., 2009. The sequence alignment/map format and samtools. bioinformatics, 25(16):2078–2079.

Linardopoulou, E. V., Williams, E. M., Fan, Y., Friedman, C., Young, J. M., and Trask, B. J., 2005. Human subtelomeres are hot spots of interchromosomal recombination and segmental duplication. Nature, 437(7055):94–100.

Liu, Q., Jiang, C., Xu, J., Zhao, M.-T., Van Bortle, K., Cheng, X., Wang, G., Chang, H. Y., Wu, J. C., and Snyder, M. P., et al., 2017. Genome-wide temporal profiling of transcriptome and open chromatin of early cardiomyocyte differentiation derived from hipscs and hescs. Circulation research, 121(4):376–391.

Logsdon, G. A., Ebert, P., Audano, P. A., Loftus, M., Porubsky, D., Ebler, J., Yilmaz, F., Hallast, P., Prodanov, T., Yoo, D., et al., 2025. Complex genetic variation in nearly complete human genomes. Nature, :1–12.

Lowe, C. B., Bejerano, G., and Haussler, D., 2007. Thousands of human mobile element fragments undergo strong purifying selection near developmental genes. Proceedings of the national academy of sciences, 104(19):8005–8010.

Malfait, J., Wan, J., and Spicuglia, S., 2023. Epromoters are new players in the regulatory landscape with potential pleiotropic roles. Bioessays, 45(10):2300012.

Mangan, R. J., Alsina, F. C., Mosti, F., Sotelo-Fonseca, J. E., Snellings, D. A., Au, E. H., Carvalho, J., Sathyan, L., Johnson, G. D., Reddy, T. E., et al., 2022. Adaptive sequence divergence forged new neurodevelopmental enhancers in humans. Cell, 185(24):4587–4603.

Marco, A., Konikoff, C., Karr, T. L., and Kumar, S., 2009. Relationship between gene co-expression and sharing of transcription factor binding sites in drosophila melanogaster. Bioinformatics, 25(19):2473–2477.

McClintock, B., 1950. The origin and behavior of mutable loci in maize. Proceedings of the National Academy of Sciences, 36(6):344–355.

Mefford, H. C. and Trask, B. J., 2002. The complex structure and dynamic evolution of human subtelomeres. Nature Reviews Genetics, 3(2):91–102.

Michalak, P., 2008. Coexpression, coregulation, and cofunctionality of neighboring genes in eukaryotic genomes. Genomics, 91(3):243–248.

Morova, T., Ding, Y., Huang, C.-C. F., Sar, F., Schwarz, T., Giambartolomei, C., Baca, S. C., Grishin, D., Hach, F., Gusev, A., et al., 2023. Optimized high-throughput screening of non-coding variants identified from genome-wide association studies. Nucleic acids research, 51(3):e18–e18.

Muerdter, F., Boryń, Ł. M., Woodfin, A. R., Neumayr, C., Rath, M., Zabidi, M. A., Pagani, M., Haberle, V., Kazmar, T., Catarino, R. R., et al., 2018. Resolving systematic errors in widely used enhancer activity assays in human cells. Nature methods, 15(2):141–149.

Muthuirulan, P., Zhao, D., Young, M., Richard, D., Liu, Z., Emami, A., Portilla, G., Hosseinzadeh, S., Cao, J., Maridas, D., et al., 2021. Joint disease-specificity at the regulatory base-pair level. Nature communications, 12(1):4161.

Neumayr, C., Pagani, M., Stark, A., and Arnold, C. D., 2019. Starr-seq and umi-starr-seq: assessing enhancer activities for genome-wide-, high-, and low-complexity candidate libraries. Current protocols in molecular biology, 128(1):e105.

Ninfali, C., Siles, L., Esteve-Codina, A., and Postigo, A., 2023. The mesodermal and myogenic specification of hescs depend on zeb1 and are inhibited by zeb2. Cell Reports, 42(10).

Numanagić, I., Gökkaya, A. S., Zhang, L., Berger, B., Alkan, C., and Hach, F., 2018. Fast characterization of segmental duplications in genome assemblies. Bioinformatics, 34(17):i706–i714.

Nurk, S., Koren, S., Rhie, A., Rautiainen, M., Bzikadze, A. V., Mikheenko, A., Vollger, M. R., Altemose, N., Uralsky, L., Gershman, A., et al., 2022. The complete sequence of a human genome. Science, 376(6588):44–53.

Nuttle, X., Giannuzzi, G., Duyzend, M. H., Schraiber, J. G., Narvaiza, I., Sudmant, P. H., Penn, O., Chiatante, G., Malig, M., Huddleston, J., et al., 2016. Emergence of a homo sapiens-specific gene family and chromosome 16p11. 2 cnv susceptibility. Nature, 536(7615):205–209.

Ohno, S., 1970. Evolution by Gene Duplication. Springer-Verlag.

Oomen, M. E. and Torres-Padilla, M.-E., 2024. Jump-starting life: Balancing transposable element co-option and genome integrity in the developing mammalian embryo. EMBO reports, 25(4):1721–1733.

Pavlovic, B. J., Fox, D., Schaefer, N. K., and Pollen, A. A., 2022. Rethinking nomenclature for interspecies cell fusions. Nature Reviews Genetics, 23(5):315–320.

Peng, T., Zhai, Y., Atlasi, Y., Ter Huurne, M., Marks, H., Stunnenberg, H. G., and Megchelenbrink, W., 2020. Starr-seq identifies active, chromatin-masked, and dormant enhancers in pluripotent mouse embryonic stem cells. Genome biology, 21(1):243.

Perry, G. H., Dominy, N. J., Claw, K. G., Lee, A. S., Fiegler, H., Redon, R., Werner, J., Villanea, F. A., Mountain, J. L., Misra, R., et al., 2007. Diet and the evolution of human amylase gene copy number variation. Nature genetics, 39(10):1256–1260.

Philipsen, S. and Hardison, R. C., 2018. Evolution of hemoglobin loci and their regulatory elements. Blood Cells, Molecules, and Diseases, 70:2–12.

Polak, P. and Domany, E., 2006. Alu elements contain many binding sites for transcription factors and may play a role in regulation of developmental processes. BMC genomics, 7:1–15.

Porubsky, D., Guitart, X., Yoo, D., Dishuck, P. C., Harvey, W. T., and Eichler, E. E., 2025. Svbyeye: A visual tool to characterize structural variation among whole-genome assemblies. Bioinformatics, 41(6):btaf332.

Razin, S. V., Ioudinkova, E. S., Kantidze, O. L., and Iarovaia, O. V., 2021. Co-regulated genes and gene clusters. Genes, 12(6):907.

Richard, D., Liu, Z., Cao, J., Kiapour, A. M., Willen, J., Yarlagadda, S., Jagoda, E., Kolachalama, V. B., Sieker, J. T., Chang, G. H., et al., 2020. Evolutionary selection and constraint on human knee chondrocyte regulation impacts osteoarthritis risk. Cell, 181(2):362–381.

Rocha, J. L., Lou, R. N., and Sudmant, P. H., 2024. Structural variation in humans and our primate kin in the era of telomere-to-telomere genomes and pangenomics. Current opinion in genetics & development, 87:102233.

Sahu, B., Hartonen, T., Pihlajamaa, P., Wei, B., Dave, K., Zhu, F., Kaasinen, E., Lidschreiber, K., Lidschreiber, M., Daub, C. O., et al., 2022. Sequence determinants of human gene regulatory elements. Nature genetics, 54(3):283–294.

Sankaran, V. G. and Orkin, S. H., 2013. The switch from fetal to adult hemoglobin. Cold Spring Harbor perspectives in medicine, 3(1):a011643.

Sanson, K. R., Hanna, R. E., Hegde, M., Donovan, K. F., Strand, C., Sullender, M. E., Vaimberg, E. W., Goodale, A., Root, D. E., Piccioni, F., et al., 2018. Optimized libraries for crispr-cas9 genetic screens with multiple modalities. Nature communications, 9(1):5416.

Sartorelli, V. and Lauberth, S. M., 2020. Enhancer rnas are an important regulatory layer of the epigenome. Nature structural & molecular biology, 27(6):521–528.

Schmidt, E. R., Kupferman, J. V., Stackmann, M., and Polleux, F., 2019. The human-specific paralogs srgap2b and srgap2c differentially modulate srgap2a-dependent synaptic development. Scientific reports, 9(1):18692.

Sedlazeck, F. J., Rescheneder, P., Smolka, M., Fang, H., Nattestad, M., Von Haeseler, A., and Schatz, M. C., 2018. Accurate detection of complex structural variations using single-molecule sequencing. Nature methods, 15(6):461–468.

Sequencing, C. and Consortium, A., 2005. Initial sequence of the chimpanzee genome and comparison with the human genome. Nature, 437(7055):69–87.

Shafin, K., Pesout, T., Lorig-Roach, R., Haukness, M., Olsen, H. E., Bosworth, C., Armstrong, J., Tigyi, K., Maurer, N., Koren, S., et al., 2020. Nanopore sequencing and the shasta toolkit enable efficient de novo assembly of eleven human genomes. Nature biotechnology, 38(9):1044–1053.

Sharp, A. J., Locke, D. P., McGrath, S. D., Cheng, Z., Bailey, J. A., Vallente, R. U., Pertz, L. M., Clark, R. A., Schwartz, S., Segraves, R., et al., 2005. Segmental duplications and copy-number variation in the human genome. The American Journal of Human Genetics, 77(1):78–88.

Smolka, M., Paulin, L. F., Grochowski, C. M., Horner, D. W., Mahmoud, M., Behera, S., Kalef-Ezra, E., Gandhi, M., Hong, K., Pehlivan, D., et al., 2024. Detection of mosaic and population-level structural variants with sniffles2. Nature biotechnology, 42(10):1571–1580.

Soler, E., Andrieu-Soler, C., De Boer, E., Bryne, J. C., Thongjuea, S., Stadhouders, R., Palstra, R.-J., Stevens, M., Kockx, C., van IJcken, W., et al., 2010. The genome-wide dynamics of the binding of ldb1 complexes during erythroid differentiation. Genes & development, 24(3):277–289.

Soto, D. C., Uribe-Salazar, J. M., Kaya, G., Valdarrago, R., Sekar, A., Haghani, N. K., Hino, K., La, G., Mariano, N. A. F., Ingamells, C., et al., 2025. Human-specific gene expansions contribute to brain evolution. Cell, 188(19):5363–5383.

Srinivasachar Badarinarayan, S. and Sauter, D., 2021. Switching sides: how endogenous retroviruses protect us from viral infections. Journal of Virology, 95(12):10–1128.

Sundaram, V., Cheng, Y., Ma, Z., Li, D., Xing, X., Edge, P., Snyder, M. P., and Wang, T., 2014. Widespread contribution of transposable elements to the innovation of gene regulatory networks. Genome research, 24(12):1963–1976.

Teichmann, S. A. and Babu, M. M., 2002. Conservation of gene co-regulation in prokaryotes and eukaryotes. TRENDS in Biotechnology, 20(10):407–410.

Thakore, P. I., D’ippolito, A. M., Song, L., Safi, A., Shivakumar, N. K., Kabadi, A. M., Reddy, T. E., Crawford, G. E., and Gersbach, C. A., 2015. Highly specific epigenome editing by crispr-cas9 repressors for silencing of distal regulatory elements. Nature methods, 12(12):1143–1149.

Thornburg, B. G., Gotea, V., and MakaŁowski, W., 2006. Transposable elements as a significant source of transcription regulating signals. Gene, 365:104–110.

Torres, D. E., Kramer, H. M., Tracanna, V., Fiorin, G. L., Cook, D. E., Seidl, M. F., and Thomma, B. P., 2024. Implications of the three-dimensional chromatin organization for genome evolution in a fungal plant pathogen. Nature Communications, 15(1):1701.

Uyehara, C. M. and Apostolou, E., 2023. 3d enhancer-promoter interactions and multi-connected hubs: Organizational principles and functional roles. Cell reports, 42(4).

Vaill, M., Kawanishi, K., Varki, N., Gagneux, P., and Varki, A., 2023. Comparative physiological anthropogeny: exploring molecular underpinnings of distinctly human phenotypes. Physiological Reviews, 103(3):2171–2229.

van der Auwera, G. and O’Connor, B., 2020. Genomics in the Cloud: Using Docker, GATK, and WDL in Terra. O’Reilly Media, Incorporated.

Vian, L., Pękowska, A., Rao, S. S., Kieffer-Kwon, K.-R., Jung, S., Baranello, L., Huang, S.-C., El Khattabi, L., Dose, M., Pruett, N., et al., 2018. The energetics and physiological impact of cohesin extrusion. Cell, 173(5):1165–1178.

Vollger, M. R., Guitart, X., Dishuck, P. C., Mercuri, L., Harvey, W. T., Gershman, A., Diekhans, M., Sulovari, A., Munson, K. M., Lewis, A. P., et al., 2022. Segmental duplications and their variation in a complete human genome. Science, 376(6588):eabj6965.

Wan, J., van Ouwerkerk, A., Mouren, J.-C., Heredia, C., Pradel, L., Ballester, B., Andrau, J.-C., and Spicuglia, S., 2025. Comprehensive mapping of genetic variation at epromoters reveals pleiotropic association with multiple disease traits. Nucleic Acids Research, 53(4):gkae1270.

Wei, X., Xiang, Y., Peters, D. T., Marius, C., Sun, T., Shan, R., Ou, J., Lin, X., Yue, F., Li, W., et al., 2022. Hicar is a robust and sensitive method to analyze open-chromatin-associated genome organization. Molecular cell, 82(6):1225–1238.

Wu, P., Li, T., Li, R., Jia, L., Zhu, P., Liu, Y., Chen, Q., Tang, D., Yu, Y., and Li, C., et al., 2017. 3d genome of multiple myeloma reveals spatial genome disorganization associated with copy number variations. Nature communications, 8(1):1937.

Xiao, F., Zhang, X., Morton, S. U., Kim, S. W., Fan, Y., Gorham, J. M., Zhang, H., Berkson, P. J., Mazumdar, N., Cao, Y., et al., 2024. Functional dissection of human cardiac enhancers and noncoding de novo variants in congenital heart disease. Nature genetics, 56(3):420–430.

Xie, K. T., Wang, G., Thompson, A. C., Wucherpfennig, J. I., Reimchen, T. E., MacColl, A. D., Schluter, D., Bell, M. A., Vasquez, K. M., and Kingsley, D. M., et al., 2019. Dna fragility in the parallel evolution of pelvic reduction in stickleback fish. Science, 363(6422):81–84.

Xie, X., Kamal, M., and Lander, E. S., 2006. A family of conserved noncoding elements derived from an ancient transposable element. Proceedings of the National Academy of Sciences, 103(31):11659–11664.

Xu, L., Liu, X., Sheng, N., Oo, K. S., Liang, J., Chionh, Y. H., Xu, J., Ye, F., Gao, Y.-G., Dedon, P. C., et al., 2017. Three distinct 3-methylcytidine (m3c) methyltransferases modify trna and mrna in mice and humans. Journal of Biological Chemistry, 292(35):14695–14703.

Yao, D., Tycko, J., Oh, J. W., Bounds, L. R., Gosai, S. J., Lataniotis, L., Mackay-Smith, A., Doughty, B. R., Gabdank, I., Schmidt, H., et al., 2024. Multicenter integrated analysis of noncoding crispri screens. Nature methods, 21(4):723–734.

Yilmaz, F., Karageorgiou, C., Kim, K., Pajic, P., Scheer, K., Consortium, H. G. S. V., Beck, C. R., Torregrossa, A.-M., Lee, C., Gokcumen, O., et al., 2024. Reconstruction of the human amylase locus reveals ancient duplications seeding modern-day variation. Science, 386(6724):eadn0609.

Yoo, D., Rhie, A., Hebbar, P., Antonacci, F., Logsdon, G. A., Solar, S. J., Antipov, D., Pickett, B. D., Safonova, Y., Montinaro, F., et al., 2025. Complete sequencing of ape genomes. Nature, :1–18.

Yu, H., Luscombe, N. M., Qian, J., and Gerstein, M., 2003. Genomic analysis of gene expression relationships in transcriptional regulatory networks. TRENDS in Genetics, 19(8):422–427.

Yu, S., Fiedler, S., Stegner, A., and Graf, W. D., 2010. Genomic profile of copy number variants on the short arm of human chromosome 8. European journal of human genetics, 18(10):1114–1120.

Yuan, X., Yan, Y., and Xue, M., 2021. Small nucleolar rna host gene 8: a rising star in the targets for cancer therapy. Biomedicine & Pharmacotherapy, 139:111622.

Zhang, D., Leng, L., Chen, C., Huang, J., Zhang, Y., Yuan, H., Ma, C., Chen, H., and Zhang, Y. E., 2022. Dosage sensitivity and exon shuffling shape the landscape of polymorphic duplicates in drosophila and humans. Nature Ecology & Evolution, 6(3):273–287.

Zhang, H., Pei, L., Ouyang, Z., Wang, H., Chen, X., Jiang, K., Huang, S., Jiang, R., Xiang, Y., and Wei, K., et al., 2023. Ap-1 activation mediates post-natal cardiomyocyte maturation. Cardiovascular research, 119(2):536–550.

Zhang, J.-Y., Roberts, H., Flores, D. S., Cutler, A. J., Brown, A. C., Whalley, J. P., Mielczarek, O., Buck, D., Lockstone, H., Xella, B., et al., 2021. Using de novo assembly to identify structural variation of eight complex immune system gene regions. PLoS computational biology, 17(8):e1009254.

Zhang, Y., Liu, T., Meyer, C. A., Eeckhoute, J., Johnson, D. S., Bernstein, B. E., Nusbaum, C., Myers, R. M., Brown, M., Li, W., et al., 2008. Model-based analysis of chip-seq (macs). Genome biology, 9:1–9.

Zheng, Y., Ay, F., and Keles, S., 2019. Generative modeling of multi-mapping reads with mhi-c advances analysis of hi-c studies. Elife, 8:e38070.

